# Quantitative Multicolored Deep Imaging of Human Bones Reveals a Composite Osteo-Sinusoidal Niche for Mesenchymal Stromal Cells

**DOI:** 10.1101/2025.10.07.680053

**Authors:** Nelson Tsz Long Chu, Ostap Dregval, Yu-Wei Chang, Emil Kriukov, Xin Tian, Xin Liu, Dana Trompet, Misty Shuo Zhang, Lei Li, Zhong Li, Emiliano Gomez Ruiz, Joana B. Pereira, Mats Brittberg, Björn Barenius, Lars Sävendahl, Ralf H Adams, Inger Gjertsson, Claes Ohlsson, Giovanni Volpe, Andrei S Chagin

## Abstract

Human bone marrow mesenchymal stromal/stem cells (BM-MSCs) are widely utilized in clinical trials and tissue engineering; however, their native microenvironment remains poorly understood. Here, we introduced a tissue-clearing protocol for human bones and integrated it with simultaneous mRNA and protein detection. Using this protocol, named DeepBone, we spatially mapped BM-MSCs relative to key bone microenvironment components, including human blood capillaries, adipocytes, sinusoids, and bony trabeculae. Quantitative analysis revealed that the native microenvironment of human BM-MSCs in young bone is enriched in vasculature, sinusoids, bone matrix, and adipocytes. In contrast, in aged bone, BM-MSCs showed no preferential association with bone or adipocytes. Proliferative BM-MSCs were predominantly found along blood vessels. Moreover, we identified a specialized microenvironment for BM-MCs in young bone, characterized by sinusoids coiled around trabeculae and enriched by R-type vessels. These findings provide novel insights into the native niches of BM-MSCs, offering a foundation for the development of tissue-engineering strategies that mimic their physiological context.

## INTRODUCTION

Mesenchymal stem cells (MSCs) in bone marrow, more accurately referred to as bone marrow mesenchymal stromal cells (BM-MSCs)^1-5^, were discovered in the 1970s as non-hematopoietic, adherent fibroblastic cells that form colonies known as fibroblastic colony-forming units (CFU) and capable of differentiating into osteoblasts, chondrocytes, and adipocytes^6^. Since the discovery of their human counterpart^7^, BM-MSCs have garnered significant interest due to their immune-modulating and regenerative potential, their ability to form chondrocytes and osteoblasts, their relatively easy accessibility via bone marrow biopsies, and their simple isolation method, often through plastic adhesion^7,8^. This interest is reflected in over fifteen thousand completed or ongoing clinical trials registered in the clinicaltrials.gov database, aimed at exploring their use in skeletal and non-skeletal regenerative medicine and cell therapies^9,10^.

Despite extensive *in vitro* characterization of human BM-MSCs (hBM-MSCs), particularly after extensive *in vitro* expansion, knowledge about naïve (uncultured) hBM-MSCs has only recently begun to emerge^1^. Recent advances in scRNAseq have begun to shed light on these cells in their naïve state^11-13^. It has been shown that hBM-MSCs are quite homogeneous, characterized by uniform expression of C-X-C motif chemokine 12 (CXCL12), platelet-derived growth factor receptors alpha and beta (PDGFRα, PDGFRβ), and leptin receptor (LepR)^11^. However, another recent report indicates some heterogeneity within human BM-MSCs, albeit with both CXCL12 and LepR still being among the most specific markers^13^.

These recent scRNAseq-based discoveries align BM-MSCs with so-called CXCL12-abundant reticular (CAR) cells^14^. These CAR cells, recently reported in human bone marrow^15^, are instrumental in supporting hematopoiesis. They form a crucial component of the stem cell niche for hematopoietic stem cells (HSCs)^16^, in part via secreting chemokines Stem Cell Factor (SCF, a product of the *Kitl* gene) and CXCL12, which regulate the retention, quiescence, and repopulation of HSCs^14,17,18^. Interestingly, when located near arterioles or sinusoids, mouse CAR cells support different subsets of HSCs^19^, pointing out the importance of the microenvironment where stem cells are situated. Furthermore, CAR cells harbor subpopulations of skeletal stem and progenitor cells (SSPCs), which give rise to cells of the osteoblast and adipocyte lineages in the bone marrow^20^.

Although CXCL12-positive BM-MSCs are critical for skeletogenesis and hematopoiesis, little is known about their native microenvironment in humans^1^. The CD146-positive subpopulation of CXCL12-positive stromal cells has been reported to localize near sinusoidal blood vessels, both within normal human bone and following bone formation by these cells in mice^21^. A recent 2D quantification of 4,704 human stromal cells against the bone surface revealed that the CD56+CXCL12+ subpopulation localizes near the bone matrix^13^. These observations underscore the limited understanding of the microenvironment inhabited by naïve human BM-MSCs^1,2^.

This knowledge gap, which hampers numerous tissue engineering and regeneration efforts, arises from the scarcity and specificity of available markers, challenges in accessing human bone tissue for modern techniques, and the complexity of bone marrow organization.

## RESULTS

### Optimizing Tissue Clearing for Aged Human Bone

To address the aforementioned knowledge gap, we first aimed to establish an optical clearing protocol for human bone that enables the detection of multiple molecular targets and is compatible with automated quantitative analysis (Fig. 1a). In parallel, we sought to integrate mRNA detection into the clearing workflow, thereby enabling spatial contextualization of single-cell RNA sequencing (scRNA-seq) data. To our knowledge, no currently available bone tissue-clearing protocols support the detection of mRNA transcripts.

**Figure 1.**
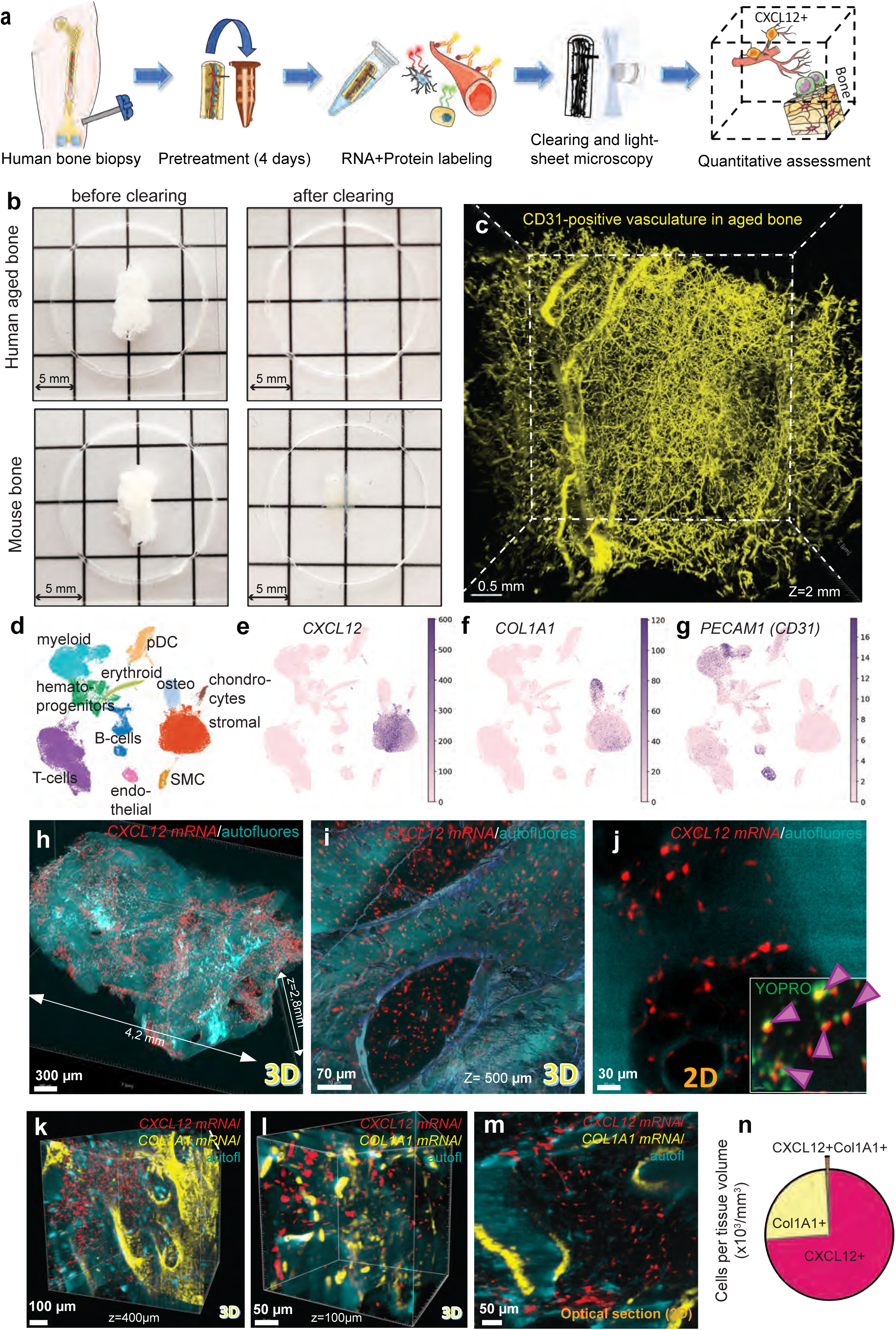
Clearing human bone and deep imaging of proteins and mRNAs. **a,** Schematic diagram of the protocol for three-dimensional characterization of human bone, including pretreatment of the bone biopsy, staining, clearing, and imaging. **b**, Images of human and murine bone samples before and after clearing. **c**, Visualization of the vasculature within human bone from a 75-year-old patient by immunostaining with antibodies against CD31. The maximum intensity projection of the overall scan with a depth (z-axis) of 2 mm. **d-g,** UMAP embedding of single-cell sequenced (scRNAseq) human nucleated bone marrow cells resulted in 11 major Leiden clusters (**d**), and feature plots demonstrating the expression of stromal cell marker *CXCL12* (**e**), osteoblast marker *COL1A1* (**f**), and endothelial cell marker CD31 (gene name *PECAM1*) (**g**) across the dataset. Cluster annotation in **d** is based on Scanpy’s rank_genes_groups and known markers. **h-j** Three-dimensional visualization of CXCL12 mRNA (pseudocolored in red) at low (**h**) and high (**i**) magnification and (**j**) representative two-dimensional (2D) optical section from the middle of the 3D scan shown in **h**. Insert in **j** shows the colocalization of CXCL12+ signals with YOPRO-1 nuclei fluorescence (pseudocolored in green) signal. YOPRO was omitted in **h-j** for better visualization. Autofluores – autofluorescence excited at 405 nm (pseudocolored in cyan). The scale bars in 3D projections should be considered approximate as they depend on the proximity to the frontal plane. The size of the aged bone sample stained and scanned in **h** is 4.2 mm (length) and 2.8 mm (depth). Purple arrow marks double CXCL12+YOPRO+ nuclei. **k-m**, Simultaneous detection of CXCL12 (red, stromal cells) and COL1A1 (yellow, osteoblasts) mRNAs at low (**k**) and at high (**l**) magnifications. **m**, Two-dimensional optical section from **k**. **n**, Pie chart showing quantification of cells positive for CXCL12, COL1A1, and double-positive from scans represented in **k**.

Based on an initial comparison of various published protocols^22-26^, considering their simplicity, reproducibility, and duration, we selected BoneClear^22^, a bone-optimized version of iDISCO^23^, as our starting point. In our hands, BoneClear enabled visualization of vasculature up to 600 µm deep in mouse bones (Extended Data (ED) Fig. 1a). The combinatorial tests of various permeabilization agents, including 3-[(3-Cholamidopropyl)dimethylammonio]-1-propanesulfonate (CHAPS), adopted from the SHANEL and Collagenase A from SUMIC protocols^24,25^, and incorporated de-lipidation with dichloromethane, as adapted from Adipo-Clear^27^, dramatically improved light penetration (Fig. 1b, ED Fig. 1b; see Methods section for details). The protocol’s compatibility with HCl- and Morse-based decalcification (ED Fig. 1b) substantially shortened processing time. Significant autofluorescence from the human bone marrow (ED Fig. 1b) hindered signal quantification and was most effectively quenched using copper (II) sulfate, consistent with previous findings^24^ (ED Fig. 1c). Blood cell decolorization in bone marrow was achieved using H₂O₂. The optimized protocol successfully visualized blood vessels in large sections of aged human bone, reaching depths of up to 3 mm (Fig. 1c and Video S1). Altogether, these initial optimizations enabled the optical clearing of human bone.

### Integrating mRNA Detection in Cleared Human Bone

Given the extensive development and informative power of single-cell RNA sequencing (scRNA-seq), we reasoned that incorporating mRNA detection into the tissue-clearing pipeline would provide a valuable tool for spatial transcriptomic analysis. To visualize mRNA transcripts in three dimensions, we employed hybridization chain reaction (HCR) RNA-FISH, a method that enables multiplexed detection of mRNA targets and has previously been applied to cleared soft tissues such as DIIFCO^28,29^.

For protocol optimization, we used mouse bone and a probe targeting *Prg4* mRNA, which is characteristically expressed in the superficial zone of articular cartilage^30^. This spatially restricted expression served as an internal control for signal specificity. Implementing HCR required omitting the conventional permeabilization step (ED Fig. 2a–d), which was feasible due to the preceding delipidation, decolorization, and decalcification steps. Among the tested decalcification methods, both Morse and EDTA preserved mRNA signal, whereas HCl resulted in signal loss (ED Fig. 2e–h). We therefore selected the Morse method for its compatibility with both mRNA and protein detection, as well as its ability to substantially reduce total protocol duration. To enable immunodetection following HCR, we introduced pH neutralization with PBS and calcium chelation with EDTA (see Methods for details).

To validate the protocol in human bone and assess mRNA probe penetration through the dense matrix, we used a probe for *SOST* mRNA, which encodes sclerostin and is expressed by bone-embedded osteocytes. *SOST* mRNA was successfully detected deep within aged human bone tissue (ED Fig. 2i, Video S2). Notably, *SOST*+ cells were localized to regions displaying a strong autofluorescent signal, corresponding to bone structures (Video S2). Illumination with a 405 nm laser revealed residual autofluorescence that mirrored bone matrix architecture (ED Fig. 2j,k). Quantification of bone volume using this autofluorescence closely matched measurements based on eosin staining (ED Fig. 2l), consistent with previous findings^31^. Furthermore, bone surfaces were clearly rendered in samples excited at 405 nm (ED Fig. 2m,n).

Importantly, HCR-generated signals remained robust throughout the bone matrix, allowing for counting of positive cells. Analysis revealed up to a 2.7-fold variation in the density of *SOST*-positive osteocytes across different trabeculae (ED Fig. 2o,p), likely reflecting local mechanical or biological heterogeneity.

We concluded that the developed protocol enables the detection of cells in human bone based on their mRNA expression.

### Transcriptional Characterization of Human Bone Marrow and Spatial Assessment of Bone Stroma

We next investigated whether the spatial localization of cell populations defined by single-cell RNA sequencing (scRNA-seq) could be inferred in human bone. Previous studies have shown that differences in sample preparation—particularly between bone marrow aspirates and enzymatically digested tissue—substantially affect the captured cellular composition, especially for non-hematopoietic populations^11,13^. To minimize such biases, we integrated our scRNA-seq dataset with several recently published datasets derived from naïve human bone marrow, including both aspirate- and enzyme-digested samples^11,32-36^. The combined dataset comprised approximately 79,000 cells, which were projected into 11 transcriptionally distinct clusters using UMAP (Fig. 1d; ED Fig. 2q-s). Of these, ∼29,000 were identified as non-hematopoietic, falling into five major clusters: endothelial cells, smooth muscle cells, osteolineage cells, chondrocytes, and stromal cells (Fig. 1d–g; ED Fig. 2r,s). Cluster annotations were based on the top 50 most cluster-specific genes (Supplementary Data File 1).

Given our research interest in skeletal biology^1-3^, we concentrated on non-hematopoietic populations, particularly the stromal cell cluster. This population is of particular interest due to its relevance in bone marrow mesenchymal stromal cell (BM-MSC)-based clinical trials and tissue engineering approaches^9^. The stromal cluster exhibited expression of canonical BM-MSC markers, including PDGFRB, THY1 (CD90), NT5E (CD73), and ENG (CD105) (ED Fig. 2r). Notably, at the mRNA level, these markers were found to be also expressed by other cell types, potentially reflecting limited correlation between mRNA and protein expression^37,38^.

Consistent with previous reports^11,13^, stromal cells were strongly enriched for transcripts characteristic of so-called CXCL12-abundant reticular (CAR) cells, including CXCL12, LEPR, KITLG (stem cell factor), and EBF3 (Fig. 1e; ED Fig. 2r,s). Of these, CXCL12 was the most robustly expressed marker: 98% of stromal cells (22,434 of 22,904) expressed CXCL12, and 80% expressed LEPR (ED Fig. 2r). CXCL12 also showed high transcript abundance, averaging 162 reads per stromal cell (Fig. 1e).

These features provided a rationale for visualizing stromal cells via CXCL12 mRNA detection using the developed HCR-based protocol. Staining for CXCL12 revealed abundant CXCL12+ cells distributed throughout the bone marrow compartment (Fig. 1h–j). Dual detection with COL1A1 mRNA, an osteoblast marker, revealed distinct spatial localization (Fig. 1k–n), underscoring the method’s ability to distinguish specific cell types in optically cleared human bone.

To further validate that the detected CXCL12+ cells represent BM-MSC/CAR cells, we combined CXCL12 HCR with LEPR protein immunodetection in both bone sections and tissue-cleared samples. Confocal and light-sheet microscopy revealed double-positive cells exhibiting extended processes (ED Fig. 3a–c), consistent with the characteristic morphology of CXCL12+ stromal cells previously described in mouse and human bone marrow^15,20,39^.

Collectively, these findings demonstrate that scRNA-seq-defined cell populations can be mapped in 3D space using the developed protocol, which we named *DeepBone*. However, in the absence of robust quantification, such spatial data remain primarily qualitative.

Terminology Note: To standardize terminology and account for the possibility that some BM-MSCs with lower CXCL12 expression may escape detection, we refer to CXCL12+ cells identified by HCR as “*CAR cells*” throughout the text below, where CAR refers to CXCL12-Abundant Reticular Cells ^14^.

### Automated Cell and Vessel Quantification

The high quality of the acquired imaging data prompted us to evaluate whether automated quantification could be achieved using widely accessible image analysis tools, such as Imaris software (Oxford Instruments Andor Inc.).

For automated cell detection, we employed the Imaris Spot function, which offers locally adaptive background subtraction, adjustable sensitivity thresholds, and customizable spot diameter settings. Despite the inherently high cellular density of bone marrow, this approach reliably identified nuclei labeled with the DNA dye YOPRO-1 (ED Fig. 3d–f; see also Supplementary Methods File). For automatic detection of CAR cells, we optimized detection parameters through iterative visual inspection of multiple optical sections to ensure comprehensive capture of labeled cells. Initial optimization was performed in parallel with YOPRO-1 staining to facilitate validation and localization of nucleated cells (ED Fig. 3g–l). Because Imaris detects spot objects in three dimensions, manual verification across adjacent z-planes was essential—spots detected in one optical section may not persist in neighboring planes (see Supplementary Methods File). Once validated, the full light-sheet dataset was processed, enabling quantification of thousands of cells. This fine-tuning combined with YOPRO-1 detection was particularly important for densely packed populations, such as osteoblasts aligned along bone surfaces (Fig. 1k–m, ED Fig. 4a–d). Co-staining with YOPRO-1 aided in calibrating spot detection across channels to ensure accurate identification of highly packed cells (ED Fig. 4a–d). Notably, the Spot detection function assigns 3D coordinates to the center of each detected spot, and all subsequent spatial measurements are based on these central coordinates.

To determine whether the visualized vasculature (Fig. 1c) could also be quantitatively assessed, we stitched image tiles using Terastitcher with non-rigid data placement^40^. Following stitching, the scans were denoised, and vessel structures were segmented in Imaris using surface and filament objects (see Methods and Supplementary Methods File). Segmentation thresholds were individually optimized for each scan to minimize background and maximize structure fidelity. This pipeline allowed quantification of vessel diameter, branching points, and terminal points (ED Fig. 4e–n). Spot detection could also be integrated with vessel segmentation (ED Fig. 4o–r), enabling analyses of spatial associations between cell types and vascular elements. In particular, combining spot diameter, sphericity, and surface volume thresholds was effective in distinguishing overlapping expression of markers shared between stromal and endothelial cells, such as ENG (see Supplementary Methods and subsequent figures).

Due to the large file sizes of high-resolution scans, we also evaluated how z-resolution affected computational load and detection accuracy. Comparative analysis of identical samples scanned at varying z-step sizes showed that resolutions below 5 μm offered no significant advantage in quantification accuracy, given the ∼3 μm light-sheet beam waist (Supplementary Methods File). Accordingly, subsequent datasets were acquired at 5 μm (or occasionally 4 μm) z-resolution to optimize the balance between detail and processing efficiency.

Quantitative analysis of the entire bone biopsy, obtained in connection with heap replacement due to fracture in elderly people, revealed 8,783 *CXCL12*+ cells per mm³ in human bone biopsy—a notably high density compared to 291 cells/mm³ of *COL1A1*+ osteoblasts in the same biopsy. This high abundance of CXCL12+ stromal cells may mirror findings in murine long bones, where tissue clearing and 3D imaging revealed stromal cell densities significantly exceeding prior estimates obtained by FACS-based methods^20,39^. Relatively minor spatial overlap was observed between the two populations (Fig. 1n), likely reflecting their partially overlapping transcriptional profiles (Fig. 1d–f; ED Fig. 2r).

These results demonstrate that automated, high-throughput quantification of transcriptionally defined cell populations in human bone processed with *DeepBone* is feasible using widely available software. The pipeline enables unbiased analysis of complex tissues, accommodates a range of staining combinations and imaging resolutions, and provides a scalable framework for in situ assessment of the human bone marrow microenvironment.

### CAR Cell Abundance and Association with Vasculature

We next sought to evaluate whether the developed imaging and analysis pipeline could detect biological differences in human bone samples. To this end, we utilized femoral biopsies from two clinical sources available to us: (i) adolescent children diagnosed with idiopathic tall stature and (ii) aged individuals undergoing hip replacement surgery due to fracture (Fig. 2a). While both sets of samples originate from the endochondral region of the femur, the precise anatomical locations are not identical. Therefore, all observed age-related differences should thus be viewed in the context of this anatomical heterogeneity and considered hypothesis-generating rather than definitive.

**Figure 2.**
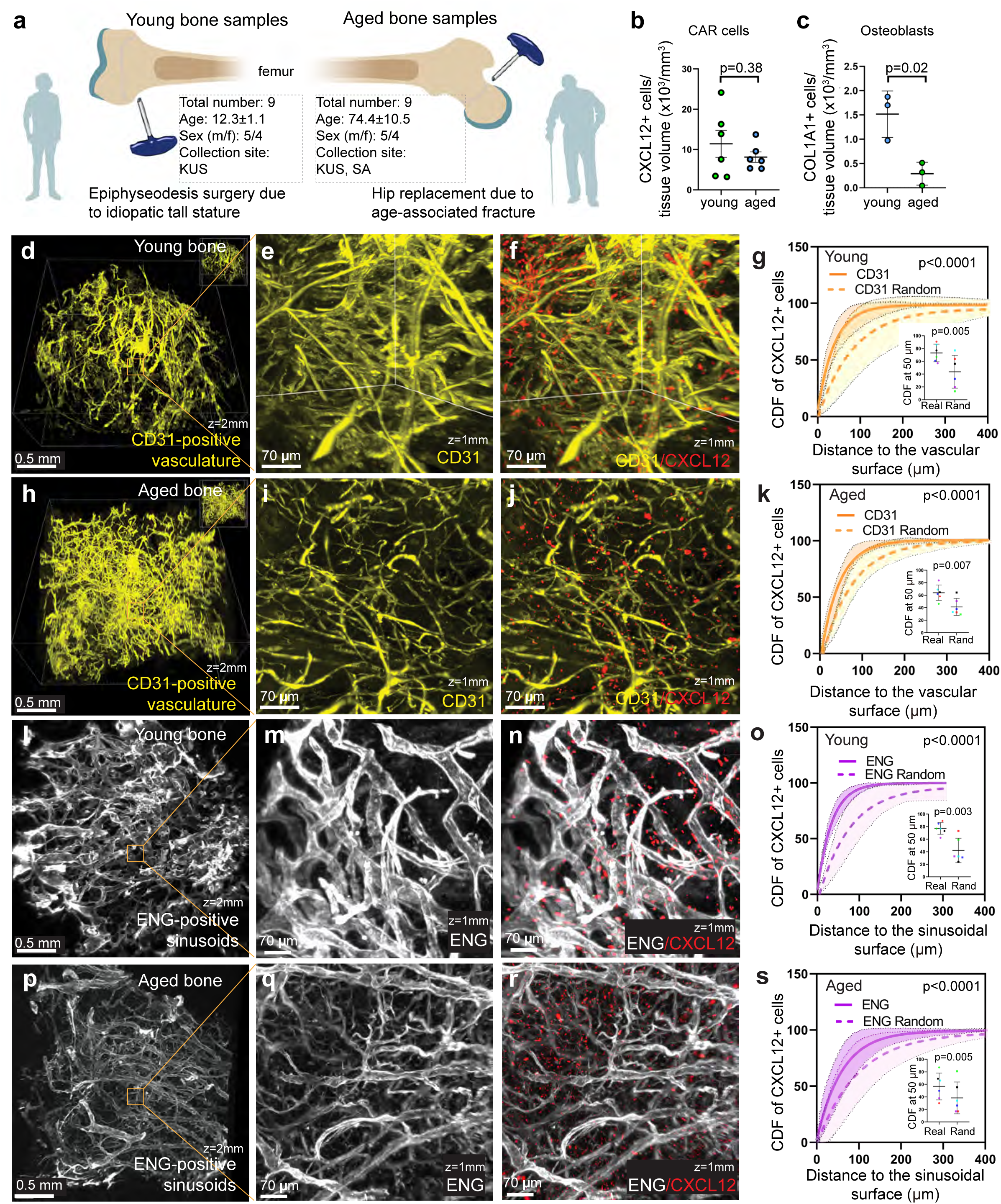
Human CAR cells are skewed toward the vasculature in both young and aged bones. **a,** Illustration depicting the collection of bone samples for comparative analysis. KUS-Karolinska University Hospital. SA-Sahlgrenska University Hospital (Sahlgrenska Academy). **b,c**, Automatic quantification of *CXCL12* (**b**) and *COL1A1* (**c**) mRNA-expressing cells in cleared bone samples obtained from young and aged patients depicted in **a**. Numbers were normalized to tissue volume. **d-k**, Three-dimensional visualization and quantification of *CXCL12* mRNA (red) together with CD31-positive vasculature (yellow) in young (**d-g,** n=6 patients) and aged (**h-k,** n=6 patients) bone samples at low (**d, h**) and high (**e, f, i, j**) magnification. The red channel is omitted in **d, e, h,** and **i**. Cumulative distribution function (CDF) of CXCL12+ cells relative to the vascular surface determined by non-linear fit with an exponential plateau (solid lines) as compared with hypothetical (random) distributions of these same spots within the same scans (dashed lines) in young (**g**) and aged (**k**) bones. **l-s**, Three-dimensional visualization and quantification of CXCL12 mRNA (red) together with endoglin (ENG)-positive sinusoids (white) in young (**l-o,** n=6) and aged (**p-s,** n=6) bone samples at low (**l, p**) and high (**m, n, q, r**) magnification. The red channel is omitted in **l, m, p,** and **q**. CDFs of CXCL12+ cells relative to the sinusoidal surface (solid lines) as compared with random distributions of these same spots within the same scans (dashed lines) in young (**o**) and aged (**s**) bones. The scale bars in 3D projections should be considered approximate as they depend on the proximity to the frontal plane. The CDFs are the mean of 6 patients (n=6), and shadow areas in **g, k, o,** and **s** outline 95% confidence intervals for these patients, and inserts depict the CDF at a 50-μm distance with data color-coded by individual patients. Real–measured data, Rand–randomized distribution. The randomized distribution of spots was performed using Imaris software. The comparison of CD functions was performed using the Kolmogorov-Smirnov test, whereas the CDF values at a 50-μm distance were compared using a paired t-test.

We first quantified CAR cells in both age groups by staining for CXCL12 mRNA. CAR cell abundance was similar between young and aged individuals (Fig. 2b), in contrast to COL1A1+ osteoblasts, which were reduced by 5.2-fold in aged bone (Fig. 2c). This reduction remained significant when normalized to bone volume: 3,783 ± 340/mm³ in young bone versus 1,437 ± 525/mm³ in aged bone (n = 3 per group; mean ± SE; p = 0.010). The CAR cell abundance and plausible preservation of their numbers with age is consistent with murine findings, where CAR cell populations are more abundant than previously estimated by FACS and appear resistant to age-related decline^20,39^.

To assess the spatial relationship between CAR cells and vasculature, we co-stained for CD31 and CXCL12 in cleared human bone (Fig. 2d-f,h-j). On average, CAR cells were more closely located to CD31+ vessels in young samples than in aged (ED Fig. 5a). Detailed spatial analysis revealed median distances of 24.6 µm in young bone and 36.9 µm in aged bone. Furthermore, 43.1% of CAR cells in young samples were located within 20 µm of a vessel surface, compared to 29.5% in aged samples (ED Fig. 5b; 57,727 cells quantified from 12 patients). A threshold distance of 20 µm (cell center to vessel surface) was used as a proxy for potential direct cell–cell interactions.

To determine whether CAR cells are non-randomly distributed in relation to blood vessels, we calculated the cumulative distribution function (CDF)—i.e., the probability of finding a CXCL12+ cell closer to the vascular surface than would be expected under a random distribution within the same scan. This analysis revealed that CAR cells were significantly enriched near blood vessels in both young and aged bone samples (Fig. 2g,k). A direct comparison between the two age groups, while acknowledging the anatomical limitations discussed above, showed that the degree of vascular proximity skewing was overall similar (ED Fig. 5c).

To test whether age-related spatial changes in CAR cells could be explained by vascular remodeling, we quantified CD31+ vessel volume. Aged bone exhibited slightly increased vascular density and total CD31+ vessel volume per tissue area (ED Fig. 5d–h). These results indicate that the marginally greater proximity of CAR cells to vasculature in young bone is not attributable to a denser vascular network.

These results demonstrate that human CAR cells are highly abundant in both young and aged bone marrow, at least seven times more numerous than osteoblasts, indicating the capacity of this approach for quantitative cell enumeration. The high abundance of CAR cells observed in human samples and their plausible age-resistance echoes findings in mouse models, suggesting a conserved spatial architecture across species. While these cells are enriched near the vasculature in both age groups, in aged patients, CAR cells were dispersed further away from the vasculature. The overall organization of the vascular bed remains relatively preserved with age and does not fully account for observed changes in spatial proximity.

### The Sinusoidal Niche for CAR cells

Beyond general vasculature, bone marrow contains sinusoidal capillaries—specialized vessels forming in hematopoietic organs. In mice, these sinusoids are well-characterized by endomucin positivity and are essential components of the adipocyte-primed niche for BM-MSCs^41-43^. In contrast, human sinusoidal capillaries have not been quantitatively assessed, likely due to the lack of good antibodies^15,21^.

To address this, we investigated whether endoglin (ENG/CD105)—previously shown to label sinusoidal capillaries in human liver^44^—could identify sinusoidal structures in human bone. Staining cleared bone samples from both age groups revealed a dense ENG+ vascular network (Fig. 2l,m,p,q). While ENG also labels non-endothelial cell types (ED Fig. 2r), these were excluded during image analysis (Supplementary Methods File). Dual staining with CD31 confirmed that ENG+ vessels were largely CD31+, although narrow CD31-bright capillaries lacked ENG staining (ED Fig. 5i–p). Quantitative analysis showed that ENG+ vessels were approximately twice the diameter of CD31+ vessels and exhibited distinct behavior when comparing samples obtained from young and aged individuals (ED Fig. 5q–u), consistent with sinusoidal identity.

We next analyzed CAR cell proximity to the ENG+ sinusoidal network using 3D segmentation and CXCL12 spot detection (Fig. 2n,r; ED Fig. 5v,w). CAR cells in aged bone were located significantly further from sinusoids (ED Fig. 5x). Median CAR-to-sinusoidal surface distance was 24.3 µm in young bone and 48.1 µm in aged bone. Additionally, 44% of CAR cells were within 20 µm of a sinusoid in young samples versus only 25% in aged (ED Fig. 5x; 78,554 cells quantified from 11 patients). CDF analysis confirmed that CAR cells were significantly enriched near sinusoids in both groups (Fig. 2o,s), but this skewing was markedly stronger in young bone (ED Fig. 5y).

These findings indicate that the developed protocol can reveal biological differences between patients (ED Figure 5a,d,h,r,u,x), and further suggest that the sinusoidal niche may undergo remodeling or functional decline with aging, even as overall CAR cell abundance remains preserved.

### Trabeculae and Adipocytes as Components of the CAR Cell Niche

Bony trabeculae and adipocytes are key structural elements of the bone marrow microenvironment. To assess their role in shaping the CAR cell niche, we visualized CXCL12 mRNA-expressing cells in combination with vasculature, bone, and perilipin (PER)-positive adipocytes in both young and aged human bone biopsies (Fig. 3; Videos S4, S5). Optical sectioning confirmed complete antibody penetration across large tissue volumes (Video S6).

**Figure 3.**
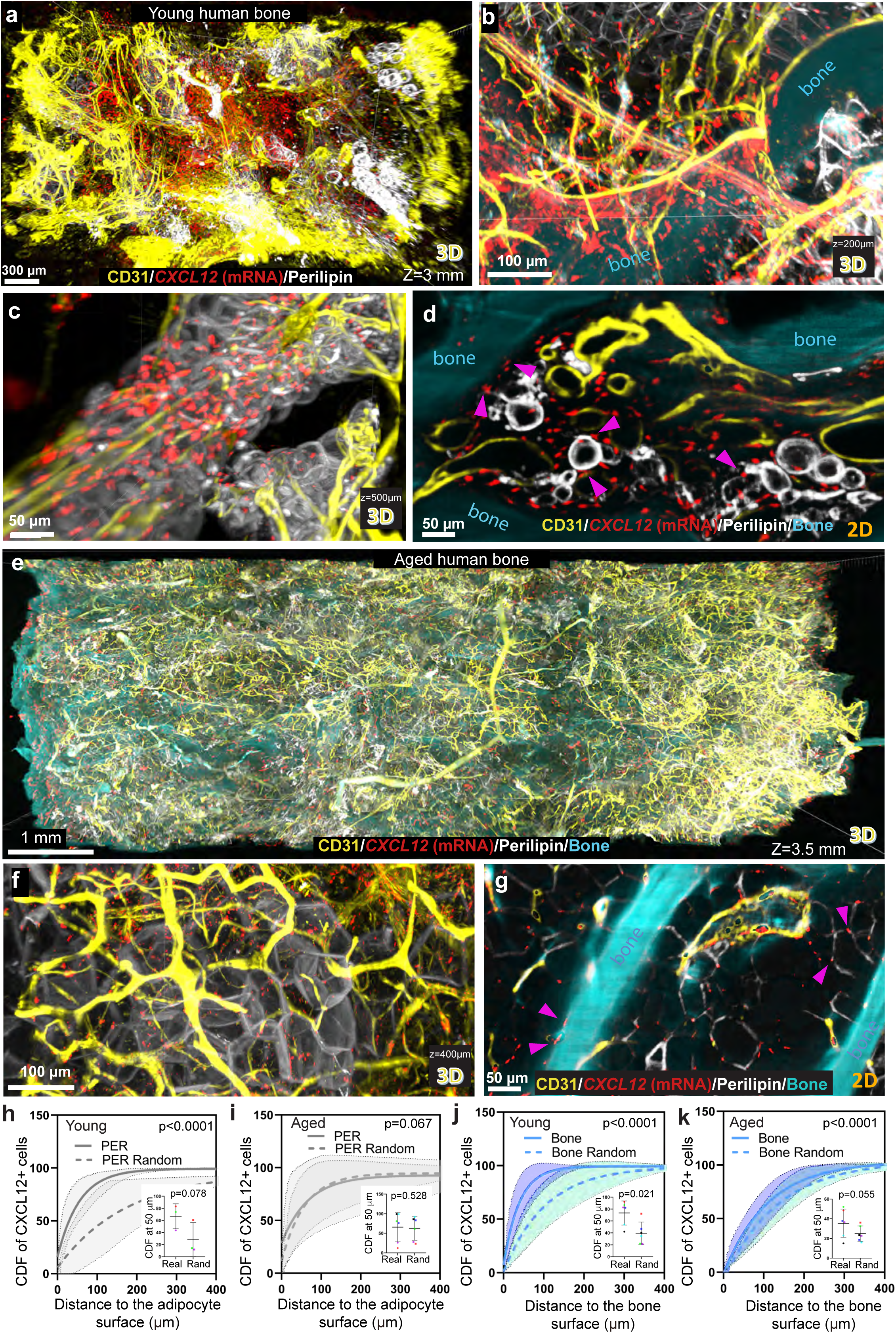
The distribution of CAR cells in relation to adipocytes and bone matrix. **a-g,** Three-dimensional *in situ* hybridization of CXCL12 mRNA (red) combined with immunostaining for CD31-positive vasculature (yellow) and perilipin-positive adipocytes (white) in young (**a-d**) and aged (**e-g**) human bone. The depth of the scans was 3 mm for young and 3.5 mm for aged bone (z-axis). Bone (cyan) was visualized on the basis of autofluorescence (omitted in **a**, **c**, and **f** to simplify visualization). **b,c,** Three-dimensional projections at a higher level of magnification, and **d** two-dimensional optical section from **a**. **f**, Three-dimensional projections at higher magnifications along with a two-dimensional optical section (**g**) from **e**. The purple arrowheads in **d** and **g** point to CXCL12+ cells on bone or adipocyte surfaces. **h-k**, Cumulative distribution functions (CDF, solid lines) of CXCL12+ cells relative to the adipocyte surface (**h**,**i,** n=3 for young patients, n=6 for aged patients) and bone surface (**j**,**k,** n=6 for young patients, n=6 for aged patients) as compared with random distributions of these same spots within the same scans (dashed lines). The shadow areas outline 95% confidence intervals for the mean (n=patients), and inserts depict the CDF at a 50-μm distance with data color-coded by individual patients. Real–measured data, Rand–randomized distribution. The randomized distribution of spots was performed using Imaris software. The comparison of CD functions was performed using the Kolmogorov-Smirnov test, whereas the CDF values at a 50-μm distance were compared using a paired t-test. A small number of patients in h,i is caused by the limited tissue availability and priorities toward other analyses.

As expected, bone adiposity increased with age (Fig. 3a–g). Interestingly, in young samples, adipocytes formed organized, gland-like clusters (Fig. 3a,c; ED Fig. 6a; Video S7). This structural heterogeneity— together with the non-uniform distribution of CAR cells and vasculature seen in large-scale scans (Figs. 1–3; ED Fig. 6a,b)—prompted us to construct 3D Gaussian kernel density maps. These revealed spatial heterogeneity on the millimeter scale in both age groups (ED Fig. 6c,d), with regions rich in CXCL12+ cells largely overlapping with highly vascularized areas. Such compartmentalization may reflect functionally distinct marrow zones, potentially linked to remodeling or hematopoiesis. Notably, clonal hematopoiesis and selective homing of multiple myeloma cells have been reported to occur in discrete marrow regions^11,45,46^, consistent with the stromal heterogeneity observed here.

In young bones, CAR cells localized near both adipocytes and trabecular bone surfaces (Fig. 3a–d; ED Fig. 7a–c). In contrast, aged bones showed more CAR cells confined between adipocytes and fewer along bone surfaces (Fig. 3f,g; ED Fig. 7d–f). Whether proximity to these compartments influences CAR cell lineage commitment toward adipogenic or osteogenic fates remains an open question^47,48^.

Spatial quantification revealed that despite the increase in total adiposity, the median distance of CAR cells to the nearest adipocyte increased with age—from 20.3 µm in young bone to 56.7 µm in aged bone. Correspondingly, 50.0% of CXCL12+ cells in young bone were found within 20 µm of adipocytes, compared to only 26.2% in aged bone (ED Fig. 7g; 94,491 cells quantified from 9 patients). CDF analysis showed a significant non-random enrichment of CAR cells near adipocytes in young bone (Fig. 3h), whereas this association was largely lost in aged bone (Fig. 3i).

Regarding the trabecular bone surface, young samples exhibited strong enrichment: 66.1% of CAR cells were located within 20 µm, with a median distance of just 10.8 µm (ED Fig. 7h). This enrichment was further supported by CDF analysis (Fig. 3j). In aged samples, the median distance increased 7.2-fold to 78.2 µm, and only 20.7% of CAR cells remained within 20 µm of the bone surface (ED Fig. 7h; 74,762 cells quantified from 10 patients). Although aged samples still showed a degree of skewing toward the bone surface, it was markedly diminished (Fig. 3k).

Thus, in young bone, CAR cells are spatially enriched near both adipocytes and trabeculae, suggesting that these compartments contribute actively to their microenvironment. In aged bones, this spatial association weakens, particularly around adipocytes, suggesting that the structural organization of the CAR cell niche may undergo functional remodeling. The observed osteo-enrichment in youth underscores the potential importance of the osteo-niche during early life stages.

### Distribution of Proliferative CAR Cells

We next investigated whether proliferating CAR cells preferentially localize near specific microenvironmental features—namely, the bone surface or vasculature. As an initial approach, we performed confocal imaging of tissue sections stained for the proliferation marker Ki67 and *CXCL12* mRNA. This validated antibody specificity and revealed heterogeneous regions of proliferative activity in the marrow. However, the number of *CXCL12*+Ki67+ cells detected was insufficient for robust quantitative analysis (ED Fig. 7i–p). This low proliferation rate is consistent with our scRNA-seq data, which showed that not many stromal cells express Ki67 mRNA (ED Fig. 2r).

Given the rarity of proliferative CAR cells, we reasoned that the broader tissue coverage and high-throughput capabilities of the *DeepBone* pipeline would enable more comprehensive quantification. To this end, we stained optically cleared human bone for *CXCL12* mRNA, Ki67, and CD31, while bone matrix was visualized via 405 nm autofluorescence (Fig. 4a–p; Video S3). Segmentation of vasculature and bone surfaces was performed (Fig. 4f,l), and single- and double-positive cells were detected in 3D and verified through optical sectioning (Fig. 4m–p).

**Figure 4.**
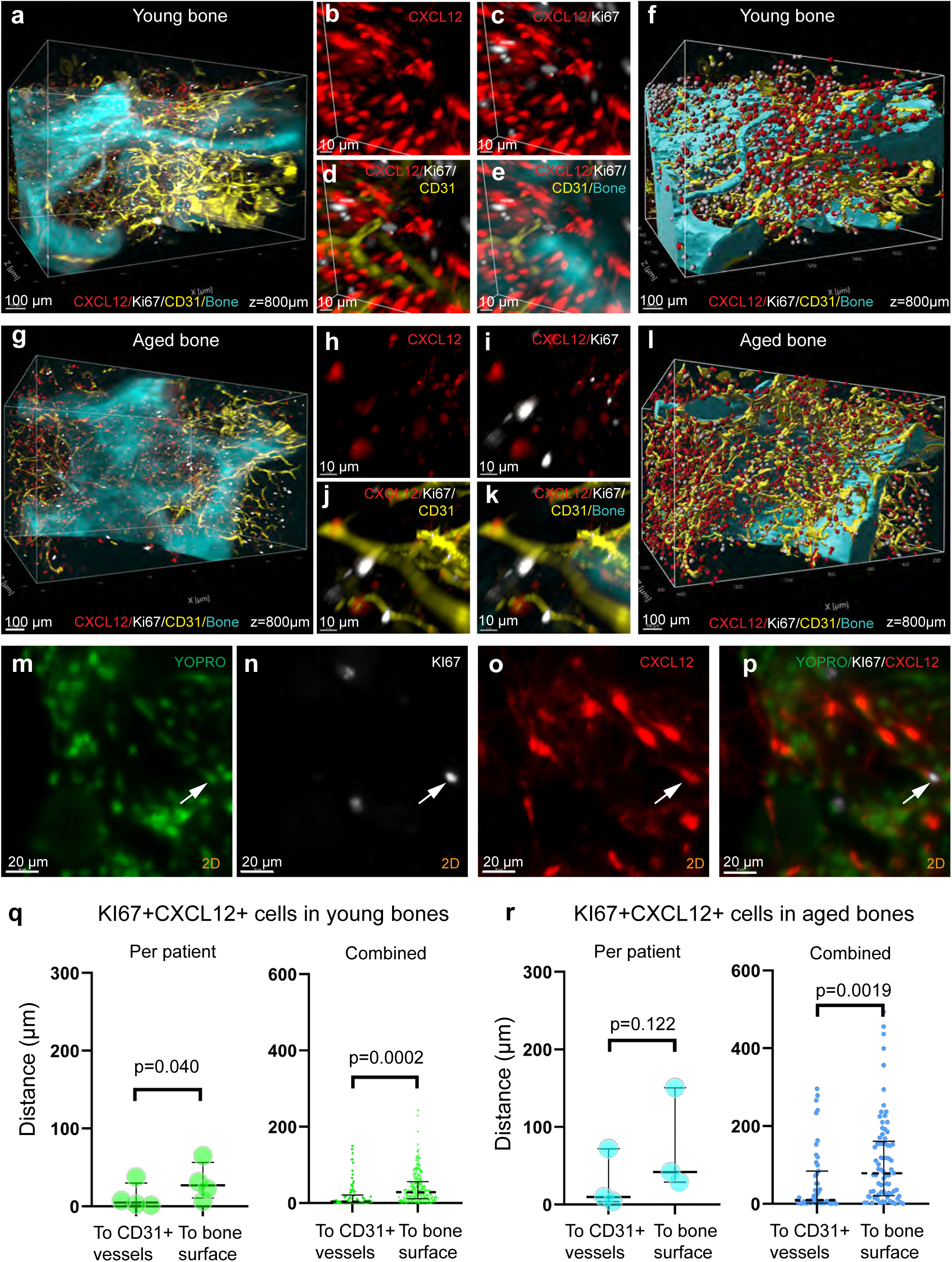
Detection and spatial characterization of dividing CAR cells in human bone. **a-l**, Multiplexed detection of proliferating cells (Ki67, white), CAR cells (CXCL12, red), blood vessels (CD31, yellow), and bone matrix (autofluorescence at 405nM, cyan) in young (**a-f**) and aged (**g-l**) human bones with original representing scans shown as a whole scanned pieces (**a,g**) and magnified volume with split channels (**b-e** from **a** and **h-k** from **g**). **f**, **l**, represent scans **a**, **g**, with positive cells spot-detected, and blood and bone surfaces segmented by Imaris software. **m-p**, Representative optical section from the light-sheet scan (**a**) with individual channels shown. The arrow depicts a KI67+CXCL12+ cell. YOPRO channel was omitted in **a-l** for visualization purposes. **q,r**, The distance from each CXCL12+Ki67+ cell to the nearest vessels and bone surfaces in young (**q**) and aged (**r**) bone samples. The data are presented as means per patient (left panels) and as violin plots (right panels) for all cells detected in these same patients to show their distribution. Dash lines in violin plots represent the median. The comparison was statistically analyzed by paired t-test (left panels, n = patients (biological replicates)) and Wilcoxon matched-pairs rank test (right panels, n = cells (individual observations)), n=4 for young patients, n=3 for aged patients.

Quantification confirmed that human CAR cells rarely proliferate. In young bone samples, only 181 *CXCL12*+Ki67+ cells were identified out of 13,275 *CXCL12*+ cells (1.36%, pooled from four donors). In aged bone, 62 proliferating CAR cells were detected among 16,223 *CXCL12*+ cells (0.38%, pooled from four donors). These rates are strikingly consistent with prior mouse data, which reported a ∼0.3% proliferation rate among stromal cells^20^.

We next examined the spatial distribution of proliferative CAR cells relative to the CD31+ vasculature and bone surface. In both age groups, proliferating CAR cells localized significantly closer to blood vessels than to bone surfaces (Fig. 4q,r). Comparative analysis between Ki67-positive and Ki67-negative CAR cells revealed a marked vascular enrichment of the proliferative subset (ED Fig. 8a,b). In young bone, 76% of Ki67+ CAR cells were found within 20 μm of a CD31+ vessel, compared to 49% of non-proliferative CAR cells; median distances were 4.9 µm and 20.5 µm, respectively (p < 0.001; ED Fig. 8a). A similar pattern was observed in aged bone (median distances: 9.9 µm vs. 36.8 µm, p<0.0001). In contrast, no significant difference was found in the proximity of Ki67+ versus Ki67-negative CAR cells to the bone surface (ED Fig. 8b).

These data indicate that under physiological conditions, human CAR cells exhibit a remarkably low proliferation rate in both young and aged bone marrow. Notably, proliferative CAR cells are consistently enriched in the immediate vicinity of the vasculature, suggesting that vascular proximity may support or regulate their proliferative capacity. This observation is consistent with the concept of osteo-angiogenic coupling reported in mouse models^43^.

### Osteo-CAR Cells Lining the Bone Surface

As shown above, a large proportion of CAR cells is enriched near the bone surface in young, but not aged, bone (Fig. 3 and ED Fig. 7). A specialized osteo-primed population of CAR cells—termed Osteo-CAR cells—has been reported in mice^48^. Given the clinical relevance of osteoporosis as an age-associated condition and our observation of a marked decline in Col1A1+ osteoblasts in aged bone (Fig. 2c), we sought to determine whether Osteo-CAR cells are reduced or spatially reorganized with age.

We first revisited our scRNA-seq dataset to assess CAR cell heterogeneity. Sub-clustering of the stromal and osteolineage populations identified five stromal sub-clusters and two osteolineage sub-clusters (Fig. 5a,b; ED Fig. 8c,d; see Supplementary File 2 for the top 50 most specific genes per cluster). All stromal sub-clusters showed high CXCL12 expression (Fig. 5b; ED Fig. 8e). Three sub-clusters (#0, #1, #2) were relatively homogeneous. Of these, sub-cluster #2 likely represents stressed cells, as indicated by elevated EGR1, FOSB, and JUNB expression (ED Fig. 8d). Based on their expression profiles, sub-cluster #3 likely reflects senescent or fibroblast-like cells, and sub-cluster #4 may represent metabolically active or stress-adaptive cells (ED Fig. 8d). The two osteolineage clusters likely represent osteoprogenitors (#5) and mature osteoblasts (#6) (Fig. 5b; ED Fig. 8d). All sub-clusters exhibited very low KI67 expression (ED Fig. 8f), supporting our imaging-based findings of limited stromal proliferation (Fig. 4).

**Figure 5.**
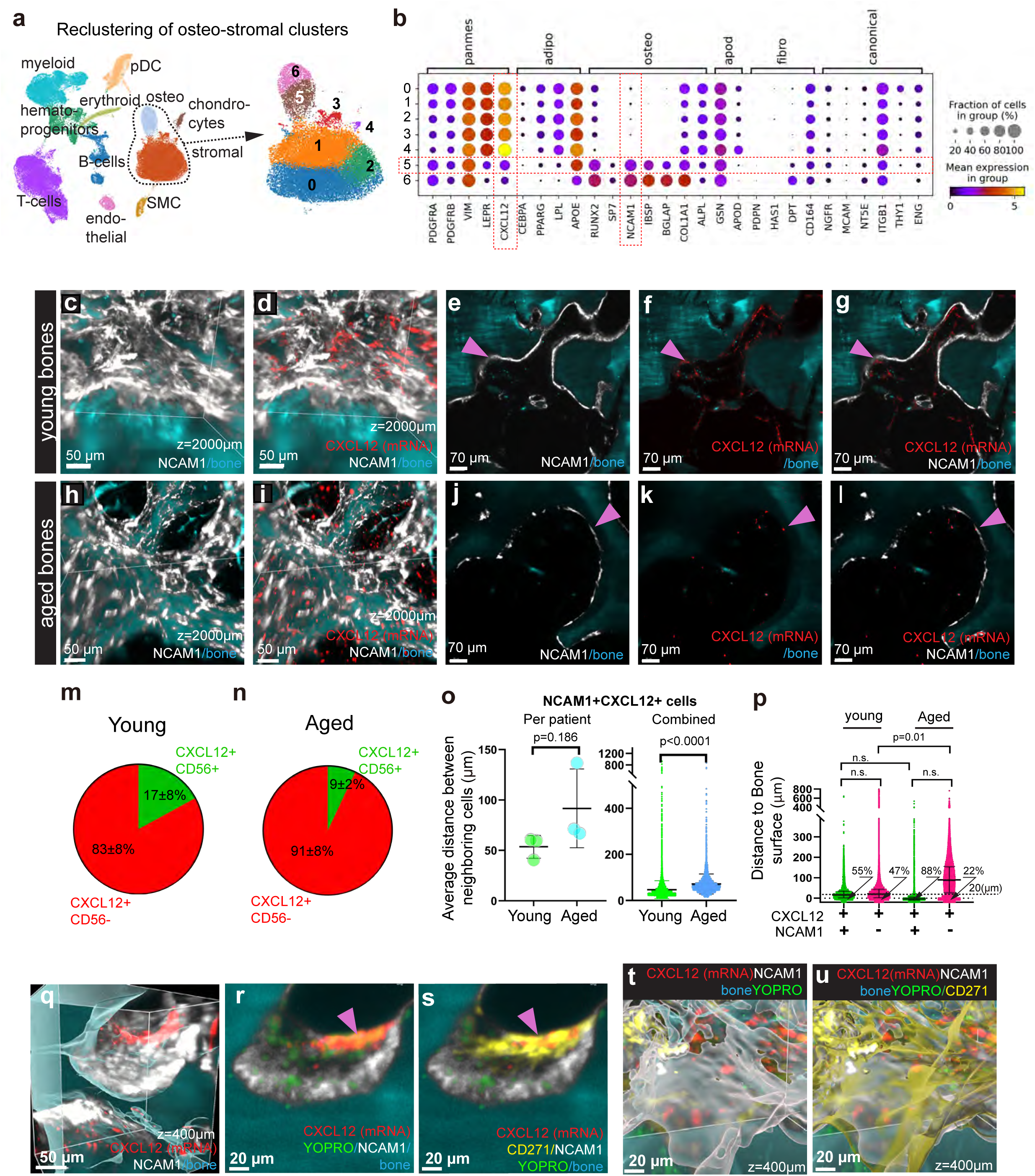
Detection and spatial characterization of osteo-progenitors. **a**, Stromal and osteo-lineage clusters were extracted from the dataset shown in Fig. 1d and re-clustered, yielding seven subclusters. **b**, Bubble plot showing expression of selected marker genes across subclusters; dot size represents the fraction of positive cells, and colour intensity denotes mean expression level. The red dashed box highlights subcluster 5, identified as osteo-progenitors, and the selected genes used for imaging. **c–l**, Multiplexed visualization of NCAM1+ (white), CXCL12+ (red) cells and bone matrix (autofluorescence at 405 nm, cyan) in young (**c–g**) and aged (**h–l**) human bone. **c,d,h,i**, Three-dimensional projections shown without (**c,h**) and with (**d,i**) CXCL12 staining. **e–g,j–l**, Optical sections from the same samples with channels separated: NCAM1 (**e,j**), CXCL12 (**f,k**), and overlay (**g,l**). Arrowheads mark double-positive CXCL12+NCAM1+ cells. **m,n**, Pie charts showing the fraction of CXCL12+NCAM1+ cells among total CAR cells in young (**m**) and aged (**n**) bone (mean ± s.d.; n = 3 patients/group). **o**, Dispersion of CXCL12+NCAM1+ cells, measured as the average distance to nine nearest neighbours. Data shown as per-patient means (left) and violin plots of pooled single-cell data (right); dashed lines indicate the median. Statistical tests: unpaired t-test (left, n = patients (biological replicates)) and Mann–Whitney U test (right, n = cells (individual observations)). **p**, Violin plots showing distances of CXCL12+NCAM1+ versus CXCL12+NCAM1-cells to the bone surface in young and aged samples (196,775 total cells analysed; 28,531 double-positive). Dashed line indicates 20 μm; percentages show the fraction of cells within this distance. One-way ANOVA was used for comparison (n = cells (individual observations)). **q–s**, Multiplexed imaging of NCAM1 (white), CXCL12 (red), CD271 (yellow), YOPRO+ nuclei (green), and bone matrix (cyan) in young bone. **q**, 3D projection with CD271 channel omitted; **r,s**, optical sections with CD271 included in **s**. Arrowheads indicate CXCL12+CD271+ cells above the NCAM1+ layer. **t,u**, 3D reconstructions with surface segmentation showing NCAM1 alone (**t**) and combined NCAM1 and CD271 (**u**); note CD271+ layer above the NCAM1+ surface.

When compared with previously reported human BM-MSC subtypes^13^, sub-cluster #0 resembles the Thy1⁺ MSC population, while clusters #1 and #2 correspond to Adipo-MSCs^13^. Sub-cluster #3 aligns with fibro-MSCs, #4 with APOD-MSCs, and #5 with Osteo-MSCs^13^ (Fig. 5b; ED Fig. 8d–i).

Based on these data, we focused on sub-cluster #5, identified previously as Osteo-MSCs¹⁴, and refer to these cells here as osteoprogenitors. In line with prior studies^13^, we used CXCL12 in combination with CD56 (NCAM1) to spatially characterize these cells. NCAM1 immunostaining revealed a distinct pattern of CD56+ cells lining the bone surface in both young and aged bone (Fig. 5c–l; ED Fig. 8j–m), consistent with its high expression in osteoblasts from our scRNA-seq data (ED Fig. 2r; ED Fig. 8i). CD56 staining was more abundant in young bone, resembling the pattern of Col1A1+ osteoblasts (Fig. 2c).

Double-positive CD56+CXCL12+ osteoprogenitors were primarily located near the bone surface (Fig. 5c–l; ED Fig. 8j–m). Quantification showed that CD56+ CAR cells accounted for 17.3 ± 8.1% of all CAR cells in young bone and 8.6 ± 1.5% in aged bone (n = 3 patients, p = 0.35, Student’s t-test; Fig. 5m,n). Supporting this decline, the mean intercellular distance (based on the nine nearest neighbors) between CD56+CXCL12+ cells was greater in aged bone (Fig. 5o). These cells localized primarily along the bone surface: in young bone, 55% of CD56+CXCL12+ cells were within 20 μm of bone, while in aged bone, this figure was 88%. Mean distances to the bone surface were 29 μm in young and 12 μm in aged bone (Fig. 5p).

In summary, CD56+CXCL12+ osteoprogenitors are predominantly found along the bone surface, and this spatial association is preserved in aged bone, despite a decline in their overall abundance.

### Osteo-niche and the Canopy Membrane

Intriguingly, in young bone, we frequently observed a dense population of CXCL12+ NCAM1-negative cells forming a layer positioned directly above the NCAM1-positive cell layer along the bone surface (Fig. 5q,r; ED Fig. 8n). Previous studies have described a specialized membranous structure in human bone, known as the canopy, that lines areas of bone formation and resorption and harbors osteogenically primed BM-MSCs^49-51^. The canopy cells have been reported to express CD271, a surface marker widely used to isolate MSCs for clinical applications^52,53^. We hypothesized that the layer of CAR cells observed above the NCAM1+ cells may represent this canopy-like structure.

To test this hypothesis, we visualized CD271+ cells in combination with CD56 and CXCL12. This analysis revealed that the CXCL12+ cells located just above the CD56+ layer were indeed CD271+ (Fig. 5q–s; ED Fig. 8n,o).

Three-dimensional segmentation of the CD56 and CD271 layers demonstrated that the CD271+ layer overlays the CD56+ layer (Fig. 5t,u). Further inspection of optical sections confirmed that the two layers are spatially distinct but often closely apposed in young bone, and consistently overlapping in aged bone (ED Fig. 8p–s). Despite the membrane-like localization and spindle-shaped morphology of CD271⁺ cells (see Video S8 and Fig. 6), NGFR (the gene encoding CD271) did not define a specific stromal subcluster in our scRNA-seq analysis (Fig. 5b; ED Fig. 8g). However, CD271+CXCL12+ double-positive cells exhibited transcriptional differences when compared to the broader stromal population (ED Fig. 8t), suggesting functional and phenotypic specificity.

**Figure 6.**
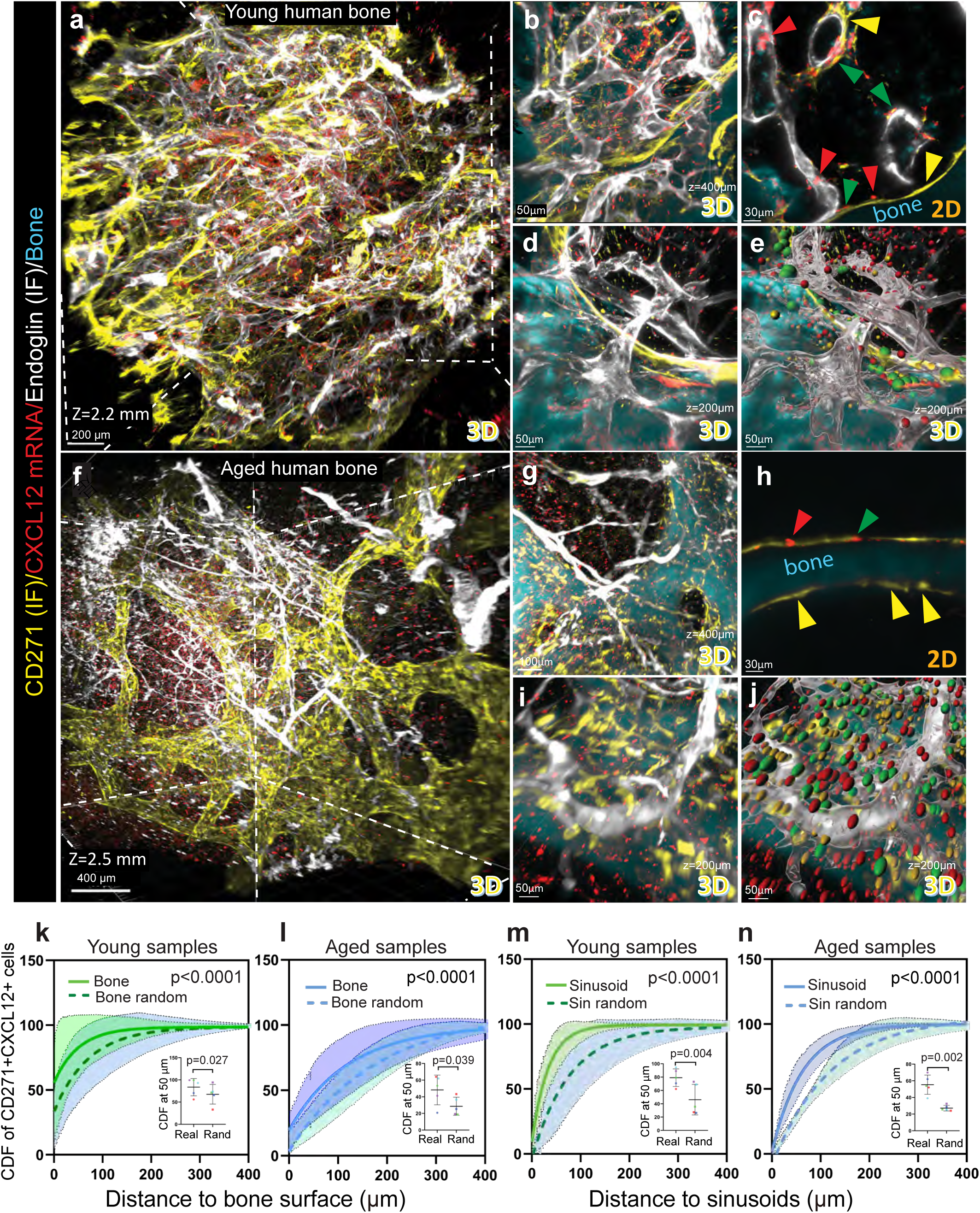
The functional CD271+ CAR cells are tightly associated with bone and sinusoids. **a-j**, Multiplexed visualization of CD271+ canopy membrane (yellow) together with CXCL12+ BM-MSCs (red), endoglin-positive vessels (white), and bone matrix (autofluorescence at 405nM, cyan) in young (**a-e**) and aged (**f-j**) human bone: **a**,**b**,**d** and **f**,**g**,**i,** three-dimensional projections at various magnifications (the Bone channel is omitted in **a** and f to improve visualization); **c** and **h**, two-dimensional optical cross-sections from **a** and **f** respectively; **e** and **j**, endoglin+ surface (grey), and spot rendering of **d** and **i** with the Imaris software, respectively. The red, yellow, and green spots (**e**,**j**) and arrowheads (**c**,**h**) indicate CXCL12+, CD271+, and double CXCL12+CD271+ cells, respectively (all detected automatically with the Imaris software). The spot size reflects the proximity of the cell to the frontal plane of the image. **l**-**n**, Cumulative distribution function (CDF) of CD271+CXCL12+ cells relative to the bone (**k**,**l**) and sinusoidal (**m**,**n**) surface in young (**k**,**m,** n=5 patients) and aged (**l**,**n,** n=5 patients) human bone. The shadow areas outline 95% confidence intervals, and inserts depict the CDF at a 50-μm distance with data color-coded by individual patients. The dashed lines depict hypothetical (random) distributions of these same spots if not associated with the corresponding surface (randomized by Imaris). Real–measured data, Rand–randomized distribution. The randomized distribution of spots was performed using Imaris software. The comparison of CD functions was performed using the Kolmogorov-Smirnov test, whereas the CDF values at a 50-μm distance were compared using a paired t-test.

To characterize CAR cells localized within the canopy membrane, we performed co-detection of CD271 and CXCL12 and analyzed the spatial distribution of double-positive (CD271+CXCL12+) cells relative to bone surfaces and sinusoids in large, cleared bone samples from both adolescents and elderly individuals (Fig. 6a–j; Video S8).

CD271+CXCL12+ cells were predominantly observed along the bone surface, with occasional positioning near sinusoidal vessels (Fig. 6c,e,h,j; ED Fig. 9a–f). Representative 3D reconstructions of double-positive cells (highlighted in green in Fig. 6e and 6j) further illustrate this spatial distribution. Notably, proximity of these cells to sinusoids appeared more frequent in young bones (Fig. 6c), suggesting potential age-related spatial remodeling of the canopy-associated CAR cell population.

Quantification revealed that a disproportionately high number of CD271+CXCL12+ cells were located within 20 µm of the bone surface, particularly in young bones — 80% in young bones and 53% in aged bones (**ED Fig. 9g,i**). Notably, in aged bones, CXCL12+ cells lacking CD271 expression were located substantially further from the bone surface, with only 17% found within 20 μm of it (**ED Fig. 9g,i**), suggesting that particularly the double-positive population makes the kinship to bone for aged stromal cells. The CDF calculation revealed that the distribution of canopy-based CAR cells was significantly skewed toward the bone surface in both ages (**Fig. 6k,l**).

Interestingly, in these analyses, we observed that sinusoids are often localized close to the bone surface, particularly in young bones (**Fig. 6b-e; ED Fig. 9a-c**). Accordingly, quantification of CD271+CXCL12+ cells in relation to sinusoids revealed that 51.9% of these cells in young bones and 36.4% in aged bones were found within 20 µm of the sinusoidal surface, with the corresponding median shortest distances 18.5 µm and 43.9 µm, respectively (**ED Fig. 9h,j**). The distribution of these double-positive cells in relation to the sinusoids was not random at both ages (**Fig. 6m,n**).

Thus, the canopy CAR cells largely remained in microenvironments highly enriched for bone and partially by sinusoids in both age groups analyzed. However, the fraction of CD271-positive CAR cells was twice smaller in aged bones, as compared with young (14.9±5.4% (n=6 aged patients) versus 28.9±3.6% (n=7 young patients), p=0.049, a total of 99,868 cells quantified; **ED Fig. 9k**). Thus, it is plausible that the sinusoid- and bone-enriched environment deteriorates with age, leading to the observed decline in the cells populating this microenvironment.

### Decoding Spatial Complexity Reveals an Osteo-Sinusoidal Niche for Human CAR Cells in Young Bones

The findings above suggest the existence of a specialized microenvironment enriched in both trabecular bone and sinusoidal vessels—a niche structure, to our knowledge, not previously described in human bone marrow. To explore this further, we analyzed the three-dimensional organization of bony trabeculae and sinusoids. 3D reconstructions revealed that sinusoidal capillaries frequently run along bone surfaces or encircle trabeculae, with CXCL12+ stromal cells interposed between these structures (**Videos S9–S11; ED Fig. 10a–d**).

Conventional spatial analysis tools are poorly suited to identifying and characterizing such complex microarchitectures. To address this, we developed a deep-learning pipeline using a graph-based variational autoencoder (VAE)—a neural network that compresses high-dimensional data into a latent representation, then reconstructs the original input while passing through an informational bottleneck (**ED Fig. 10e; see Methods for details**).

To prepare the input data, we converted multichannel 3D images into multilayer graphs, where each layer represented a distinct structure—bone, sinusoids, or CXCL12+ cells. For each CAR cell, we computed the number of nearby structures contained within spherical shells of increasing radii (0– 200 µm, 10 µm increment), yielding three radial distribution functions (bone, sinusoids, other CAR cells) that quantified the local environment (**Fig. 7a**). These functions (63 total values per cell – 3 functions multiplied by 21 radius points) served as inputs for the VAE, which compressed them into a 2D latent space and then reconstructed the original functions (**Fig. 7b; ED Fig. 10e**). In this latent space, each CAR cell is positioned according to the similarity of its microenvironment, allowing unsupervised grouping of spatially related cells.

**Figure 7.**
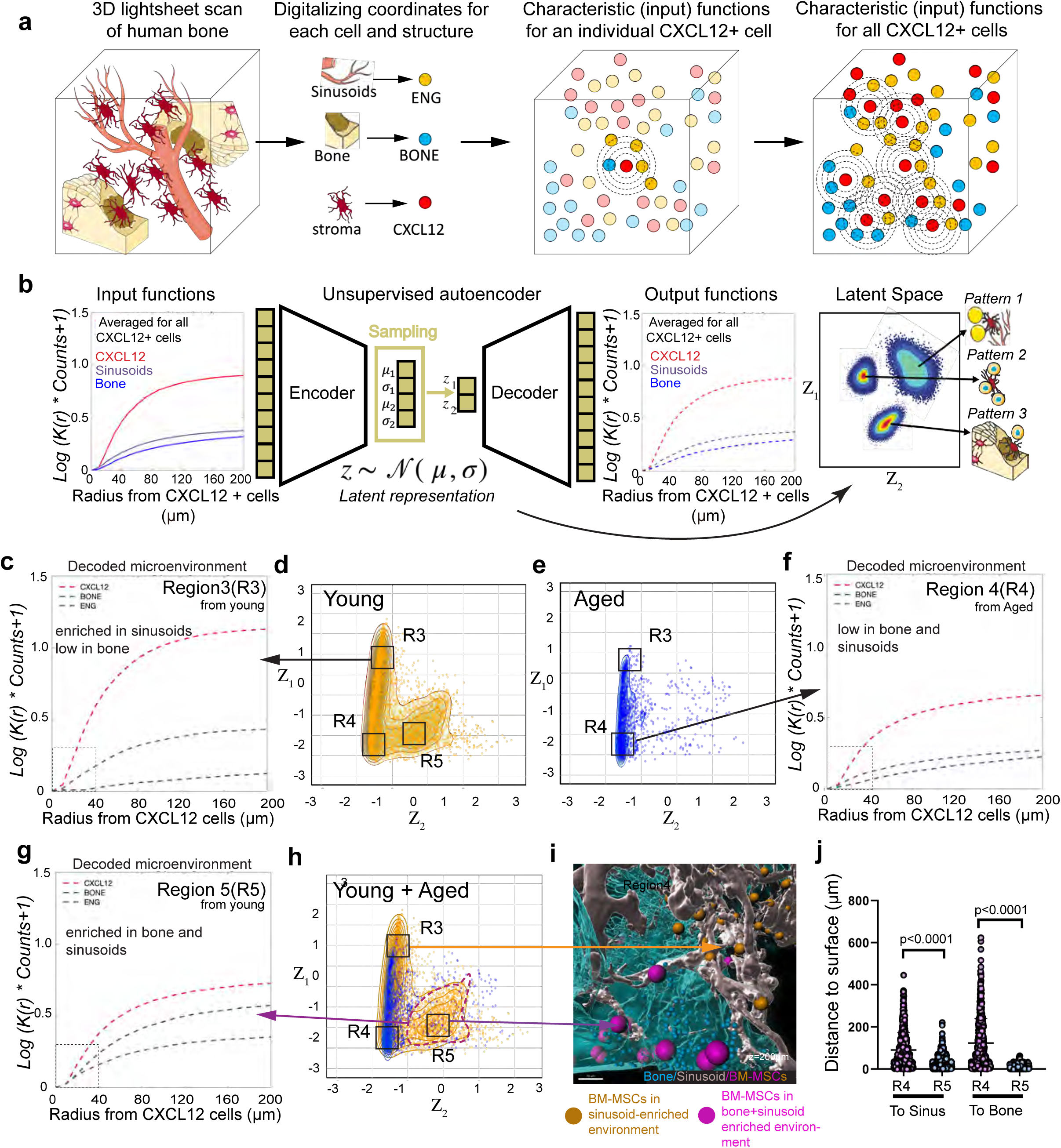
Unsupervised learning reveals an osteo-sinusoidal niche for CAR cells in young but not aged bones. **a**, A schematic diagram illustrating the steps taken to construct functions centered on CXCL12+ cells and characterizing their microenvironment. Lightsheet microscopy signals from each channel were converted to spot objects with spatial coordinates in different classes with Imaris. A sphere of varying diameter was constructed around each CXCL12+ cell for the class of CXCL12+ (red spots), bone (blue spots), and sinusoids (ENG, yellow spots). **b**, The radial input functions obtained in **a** were used to train a variational autoencoder neural network to encode all the obtained values (totaling 63) into a two-dimensional latent space (coordinates z_1_ and z_2_), and then decode these two values back to the original functions. The latent space reveals classes of micro-patterns in the 3D space surrounding the CXCL12+ cells. **c-h**, Visualization of the encoded (**d, e, h**) and decoded (**c, f, g**) environment in a latent space for CXCL12+ cells in young (**c, d**), aged (**e, f**) patients and overlap (**h**) (3 individuals per group). The decoding was done to specific regions depicted as R3, R4, and R5 in **d,h**. The decoded microenvironment is presented as cumulative curve plots of enrichment for other CXCL12+ cells, bone, and sinusoids in relation to the distance from each CXCL12+ cell within the selected region: (**c)** represent microenvironments enriched in sinusoids but low in bone, (**f)** represent microenvironments low in both sinusoids and bone, and **(g)** represents a microenvironment enriched in both bone and sinusoids. **i**, The original coordinates for CXCL12+ cells (purple dotted line) from selected regions were retrieved and imported back to the original scans (using Imaris) to visualize the local environment of the cells (**h**, young bone). **j**, Violin plot shows shortest distances of CXCL12+ cells from regions R4 and R5 (**h**) to sinusoids (sinus) and bone surfaces. Individual cells were pooled from three independent young patient samples. Solid lines represent the Mean with SD.

This approach revealed two distinct clusters of CXCL12+ cells in young bone, but only one in aged bone (**Fig. 7d,e**). Overlaying coordinates showed that the aged cluster aligned with the left-hand cluster in young bone, while the right-hand cluster was largely absent in aged samples (**Fig. 7d,e,h**). Of all cells represented in the latent space, 61.9% were unique to young bone, with only 31.2% overlapping between age groups. This suggests the presence of an age-restricted microenvironment for CAR cells in young individuals.

To further dissect this, we extracted cells from three regions of the latent space: R3 and R4 (age-stable clusters), and R5 (age-sensitive cluster) (**Fig. 7c–h**). Decoding their spatial neighborhoods revealed that R5 cells were enriched in proximity to both sinusoidal vessels and trabecular bone—defining an osteosinousoidal microenvironment. In contrast, R3 and R4 cells represented a gradient of sinusoidal proximity with minimal bone association (**Fig. 7f–g**).

To validate this results, we imported the coordinates of the CXCL12+ cells (R4, R5) identified within distinct patterns back to the original scans, which visually confirmed the presence of two distinct, spatially localized CAR niches: (i) a sinusoidal niche, present in both age groups (CAR cells in orange; **Fig. 7i; ED Fig. 10f**), and (ii) an osteo-sinousoidal niche, observed largerly in young bone (CAR cells in pink; **Fig. 7i; ED Fig. 10g**).

To further validate these niche identities, we measured distances from imported R4 and R5 CAR cells to the nearest bone and sinusoidal surface within the light-sheet scans (**Fig. 7j**). The analysis confirmed that the two latent-space-defined populations were significantly different in their proximity to these structural elements.

Thus, by applying a graph-based variational autoencoder to high-resolution spatial data, we achieved an unsupervised, unbiased quantification of CAR cell microenvironments in human bone. This approach uncovered two major stromal niches: a stable sinusoidal niche conserved across ages, and an osteo-sinousoidal niche exclusive to young bone.

### Identification of R-Type Capillaries in the Osteo-Sinusoidal Niche

Some of the sinusoidal capillaries spanning the bone surface or wrapping around trabeculae closely resemble post-arterial R-type capillaries, which were recently discovered and characterized in mice^54^. To determine whether R-type capillaries are present within the human osteo-sinusoidal niche, we first sought to identify these cells within our previously described single-cell RNA sequencing (scRNA-seq) atlas of human bone marrow. Given the relatively low number of endothelial cells (ECs) in our dataset (1,835 cells), we integrated it with a recently published dataset comprising 4,099 sequenced human bone marrow ECs^13^. UMAP clustering of the combined dataset revealed six distinct endothelial subclusters (Fig. 8a–c; ED Fig.11a-g). However, expression of canonical murine R-type vessel markers did not clearly correspond to any single human cluster (Fig. 8b).

**Figure 8.**
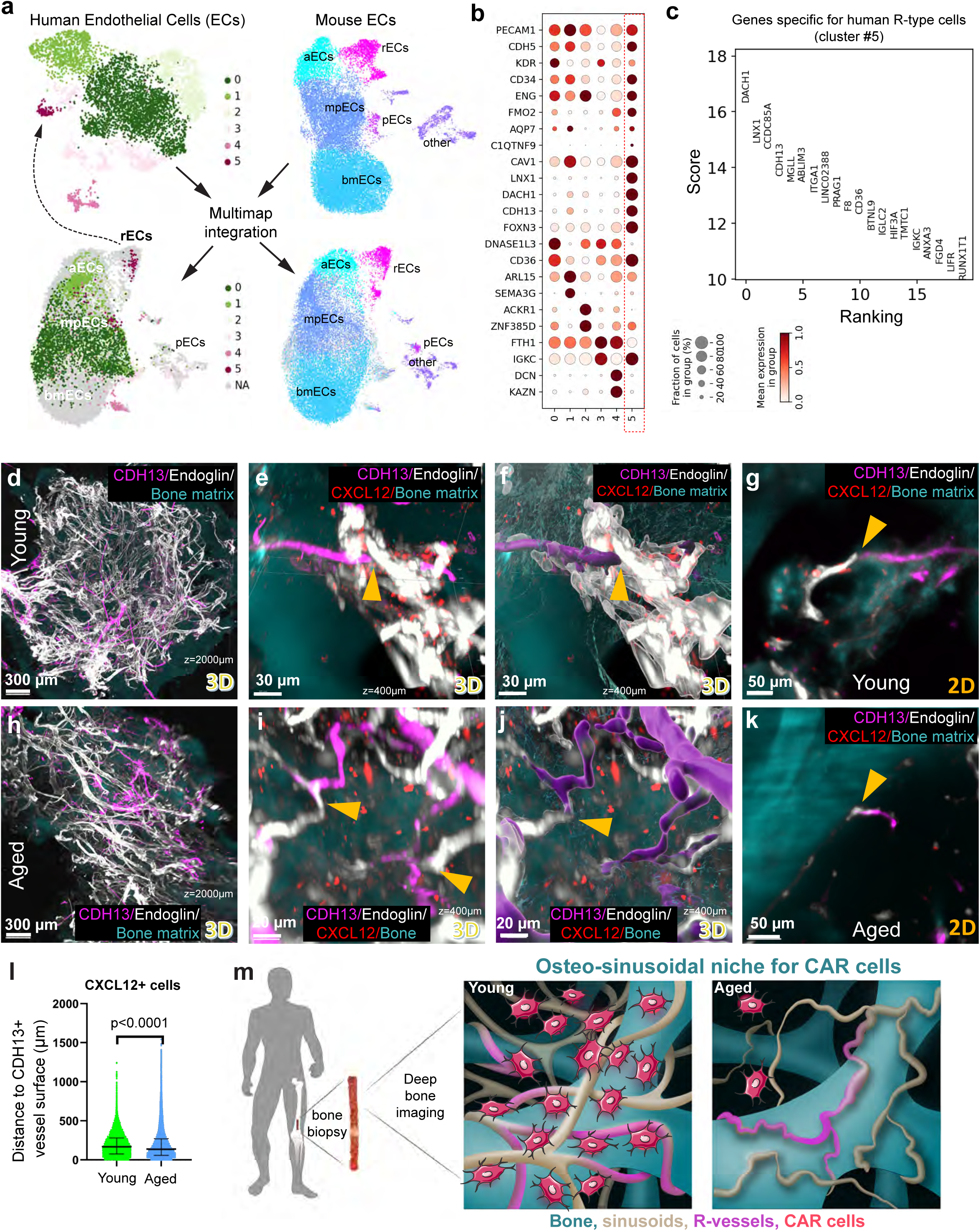
The osteo-sinusoidal niche contains R-type post-arterial vessels. **a**, MultiMAP-based integration and clustering of human endothelial cells (ECs, left panels) and mouse ECs (right panels), used to project mouse-defined clusters onto the human dataset. rECs, R-type ECs; aECs, arterial ECs; mpECs, metaphyseal ECs; bmECs, bone marrow ECs; pECs, proliferating ECs. **b**, Bubble plot showing expression of selected marker genes across human EC subclusters. Dot size indicates the proportion of positive cells, and colour intensity reflects the mean expression level. The red dashed box marks subcluster 5, identified as R-type ECs. **c**, The top-ranked genes specifically enriched in the human rEC subcluster. **d–k**, Multiplexed 3D visualization of CDH13+ R-type vessels (purple) and endoglin+ vessels (white), together with CXCL12+ cells (red) and bone matrix (cyan; autofluorescence at 405 nm), in young (**d– g**) and aged (**h–k**) human bone at low (**d,h**) and higher (**e,f,i,j**) magnification. The CXCL12 channel (red) is omitted in **d,h** to enhance vessel visualization. Panels **f,j** correspond to **e,i** and show segmented endoglin+ (grey), bone (cyan), and CDH13+ (purple) structures rendered with Imaris software. **g,k**, optical sections derived from **e,i**, respectively; orange arrowheads denote apparent junctions between R-type vessels and sinusoids. **l**, Violin plots showing the distance between CXCL12+ cells and the CDH13+ surface in young and aged human bone (n = 103,090 cells (individual observations) derived from 3 patients per group). Dashed lines indicate medians. Statistical comparison: Mann–Whitney U test. **m**, Schematic representation of the osteo-vascular niche for CAR cells in young and aged human bone.

To improve resolution and establish cross-species correspondence, we applied label transfer between mouse and human datasets (see details in the Supplementary Method File). First, we downloaded and re-clustered a published mouse endothelial cell dataset in which R-type endothelial cells had been defined^54^. The R-type cluster was readily identified based on its reported marker profile (Fig. 8a; ED Fig. 11h–l). We then performed label transfer from the mouse to the human EC dataset. While a standard PCA-based pipeline produced suboptimal alignment, two advanced methods - scANVI^55^ and MultiMap^56^ - yielded consistent results. The final analysis was based on MultiMap, which robustly aligned one human endothelial cluster (cluster 5) with the mouse R-type population (Fig. 8a).

We next examined the top 20 genes enriched in cluster 5 to assess their specificity and expression levels across other cell types (Fig. 8b, c). Based on this, we selected several candidates, including DACH1, ADGRL4, LNX1, CDH13, and FOXN3, and tested their specificity on vessel labeling. Among these, DACH1 and CDH13 emerged as particularly promising, based on their restricted expression patterns and the anatomical and morphological characteristics of the labeled vessels (ED Fig. 11m-v). As high-quality antibodies were available for CDH13, we selected this marker for further in situ validation and analysis.

Visualization revealed that many CDH13+ vessels extended along the surface of bony trabeculae (Fig. 8d–k, ED Fig. 11s,t), as would be expected from the mouse study^54^. Quantitative analysis of these vessels showed no statistically significant differences between young and aged bones (ED Fig. 11u,v; the total vessel length was 1.6 ± 0.09 × 10³ µm/mm³ in young (n=3) versus 0.7 ± 0.35 × 10³ µm/mm³ in aged samples (n=4), mean ± SE; p = 0.095). This finding contrasts with reports of age-associated increases in R-type vessel abundance in mice^54^. The discrepancy may reflect species-specific differences, as aged human bone marrow is characterized by increased adiposity and trabecular density, features that could influence vascular organization. Alternatively, our comparison may be confounded by anatomical heterogeneity across samples, as discussed above, and a more spatially standardized analysis will be needed to draw definitive conclusions.

Interestingly, some CDH13+ vessels formed self-looping structures (ED Fig. 11y,z,ac) - a hallmark of capillary networks - suggesting a role in localized oxygen and nutrient delivery, as was shown in mice^54^.

Manual inspection of 3D scans further revealed direct transitions between CDH13+ vessels and ENG+ sinusoids near the bone surface (Fig. 8d–k), indicating a structural continuum between post-arterial capillaries and sinusoidal vasculature. While current software limitations precluded direct, unbiased quantification of vessel-to-vessel junctions, as a proxy, we used filament tracing of CDH13+ vessels to identify terminal points and then calculated their distances to ENG+ sinusoids and to the bone surface in 3D space (ED Fig. 11w–ad). This analysis revealed that a substantial proportion of CDH13+ terminal points were located within 40 μm distance to both ENG+ vasculature and bone: 58.2% ± 11% in young and 30.0% ± 15% in aged bones (n=3 per group) (ED Fig. 11w,x,aa–ad).

In relation to CAR cells, spatial quantification showed that CXCL12+ stromal cells in aged bones were located slightly closer to CDH13+ vessels than in young samples (median distance: 171.1 µm in young vs. 139.4 µm in aged; n=3 per group; 103,090 cells analyzed; Fig. 8l). However, only a small fraction were found within 20 µm of CDH13+ vessels—9.4% in young and 4.8% in aged—suggesting limited direct interaction between CAR cells and R-type vessels.

Taken together, based on marker expression, anatomical distribution, and 3D reconstructions, we conclude that the osteo-sinusoidal composites observed in young bones include a distinct population of post-arterial capillaries—referred to here as R-type vessels. While their role in supporting CAR cells remains unresolved, the discovery of transition points between R-type vessels and sinusoids near the bone surface further supports the notion that the osteo-sinusoidal compartment represents an anatomically and functionally distinct microenvironment compared to the classical medullary sinusoidal niche for CAR cells^20,39^.

## DISCUSSION

This is the first 3-dimensional quantitative characterization of the microenvironment in which human CXCL12-positive BM-MSCs (CAR cells) are located in the bones of young and aged humans. We have also identified osteo-sinusoidal composites enriched by novel R-type vessels and shown that these structures form a specific niche for CAR cells in young human bones (**Fig 8m**).

Despite significant recent progress in 3D imaging of human bone, challenges associated with clearing human bone have resulted in a shortage of reports in this direction^25,26,57,58^. Here, we developed a simple, rapid, scalable, reproducible, and inexpensive protocol named *DeepBone* that allows detailed 3D characterization of the spatial organization of human bone marrow by simultaneous detection of several proteins and mRNA transcripts. The *DeepBone* protocol represents a significant advance in this context, allowing for novel analyses and insights. For example, the high quality of the data acquired with *DeepBone* permitted us to deploy new graph-theory-based and AI-powered tools for microenvironmental assessment, such as BRAPH^59^.

The possibility of detecting mRNA by *DeepBone* allowed us to integrate recent advances in single-cell RNA sequencing (scRNAseq) into a spatial context of the human BM-MSCs microenvironment. Our results reveal that human BM-MSCs/CAR cells are more abundant than previously anticipated based on fluorescence-activated cell sorting (FACS) analysis^7,60^, which aligns well with recent microscopic observations in both human and animal studies^13,20,39^. Similar to mouse data, the number of human BM-MSCs/CAR cells was relatively stable with age, and these cells populated both peri-arteriolar and peri-sinusoidal niches regardless of age. The observed very low level of proliferation of human CAR cells was remarkably similar to the one reported for mouse BM-MSCs^20^. These observations highlight similarities between murine and human stromal cells, justifying the extrapolation of numerous mouse observations to human pathophysiology. For example, in mice, CXCL12-expressing stromal cells located near sinusoids form a niche that facilitates hematopoietic stem cell (HSC) trafficking in and out of the bone marrow, whereas stromal cells near arterioles form a niche for quiescent HSCs^19^. It appears plausible that the human CAR cells observed in association with arterioles and sinusoids also provide distinct niches for HSCs.

We found that adipocytes and the bone surface play an active role in forming a microenvironment for CAR cells in an age-dependent manner. This may be attributed to substantial differences between human and mouse bones in terms of bone adiposity and trabeculation, with both parameters being more pronounced in humans. The role of adipocytes in regulating skeletogenesis is well known^61-63^, supporting their potential role in the niche composition for human CAR cells. However, our approach is limited regarding adipocytes due to the lack of scRNAseq data and the possibility that some adipocytes may express CXCL12^47^. Therefore, we cannot currently elaborate on the discrepancy between enriched adiposity and the loss of adipose-associated distribution of CAR cells with age. Nonetheless, it is noteworthy that in young bones, adipocytes often appeared in gland-like clusters, reminiscent of the recently described lobe-like architecture of human visceral adipose tissue^64^. This localized adipogenesis during skeletal growth may indicate clonal expansion of adipocyte progenitors. Moreover, it suggests that specific skeletal stem cells may be primed toward adipogenic differentiation already during early developmental stages in humans.

The location of some human stromal cells near the bone surface was recently reported^13^, strengthening our efforts to explore this niche in a three-dimensional and quantitative manner. The remarkable enrichment of bone-associated CAR cells in young bones suggests the importance of this structure in the regulation of stromal cells, at least at this age. It is reasonable to assume that CAR cells situated near the bone surface are more prone to generate osteoblasts on that same surface. Supporting this reasoning, further enrichment in localization toward the bone surface was observed among stromal cells within the canopy membrane, which lines the bone and harbors osteoprogenitors^50^. Interestingly, this canopy membrane consists of CD271-positive cells, a common marker for isolating human BM-MSCs, including in numerous clinical trials^49^.

Furthermore, 3D deep imaging revealed that sinusoids and bony trabeculae form a complex spatial network in which blood vessels run along or wrap around trabeculae, creating osteovascular composites—an architecture that cannot be appreciated without volumetric imaging. Moreover, a part of these osteo-vascular composites is formed by recently discovered R-type vessels^54^. In mice, the bone marrow has been shown to possess an extensive sinusoidal vasculature, with most stromal cells reported to reside near sinusoids^20,39,60,63,65^. However, to our knowledge, stromal cell enrichment near bone surfaces has not been reported in mice, and visual inspection of the referenced studies does not reveal any such spatial pattern. In contrast, our human bone samples, particularly from young individuals, exhibited a clear enrichment of stromal cells adjacent to bone surfaces, with this pattern being retained in aged bone for CD271+ and CD56+ subpopulations of CXCL12+ stroma. These findings suggest that the osteosinusal niche we describe may be specific to human bone marrow. This interpretation aligns with known species differences: human bone shows greater trabecular complexity, higher adiposity, and distinct vascular organization compared to mouse bone.

Our autoencoder-powered analysis revealed that these osteo-vascular composites form a unique niche for human CAR cells in bones from young but not aged individuals. Niches largely determine the balance of different progeny generated from stem cells^19,66^. Therefore, it is plausible that the age-associated decline in osteogenesis is caused not only by the intrinsic aging of the stem cells themselves^67^, but also by changes in the specialized niches observed herein.

It is important to note that, although both young and aged samples were obtained from the subchondral region of the femoral epiphysis, they are not anatomically identical. Consequently, the possibility that the differences observed between adolescent and aged patients could be due to anatomical variations, rather than aging alone, cannot be fully excluded. Therefore, age-related changes should be interpreted with caution, recognizing that anatomical differences may contribute to the findings. Additionally, the human samples were not stratified by sex and were limited in number, allowing the detection of only very pronounced and generalized differences. The developed protocol enables further research in this area.

### Outlook

Altogether, our findings emphasize the power of the 3D approach, which enabled the identification of an osteovascular niche for stromal cells, a previously underappreciated aspect of human bone marrow architecture that appears to be prominent in young but not aged bones.

These data shed light on the cellular arrangements within the bone marrow of young and aged human bones, potentially advancing our understanding of diseases such as osteoporosis, non-union fractures, and cancer metastases, while also opening new avenues for tissue engineering approaches utilizing MSCs.

### Methods

#### Ethical aspects of procedures involving human tissue

Femoral bone biopsies were collected from the subchondral region within 2 cm of cartilage. Aged human bone was retrieved from 12 aged patients, between 53 and 89 years of age, undergoing femoral head replacement surgery in association with hip fracture. Among them, three patients were collected at Karolinska University Hospital, Stockholm, Sweden, and six were collected at Sahlgrenska University Hospital, Gothenburg, Sweden. Human adolescent bone was collected from 10 patients between 11 and 14 years of age, undergoing epiphysiodesis surgery at Karolinska University Hospital, Stockholm, Sweden, in connection with predicted idiopathic tall stature. Detailed patients’ information can be found in Supplementary Table 1. Oral and written consent was obtained from all the patients and, in the case of teenagers, from their parents. The collection of human material was approved by the Swedish Ethical Research Authority and the National Board of Health and Welfare (ethical permissions numbers 2022-01977-02, 2014/276-31/2 and 97-214). Bone was collected with a bone marrow biopsy needle, fixed in 4% PFA/1X PBS for 1-2 days at 4 degrees Celsius, and further processed for 3D RNA and protein probing.

#### DeepBone protocol

The DeepBone protocol is tailored and scalable for common human bone samples collected from bone needle biopsies, but has also been tested successfully on mouse bones and soft tissues: liver, muscle, and kidney. The protocol consists of the four steps below: (i) tissue pre-treatment, (ii) 3D RNA detection, (iii) 3D immunostaining, and (iv) tissue clearing. The second or third steps can be omitted depending on detection needs. Incubation times below are provided for human bone samples ranging from 2 mm to 15 mm in length and 5 mm in diameter. These times can be shortened for smaller samples.

##### (i) Tissue Pretreatment

Harvested human bone biopsies were first washed with 1x PBS/0.5%Tween-20 (Sigma Aldrich) for 1 hour and dehydrated with consecutive methanol (Histolab Products AB) gradients from 20%, 40%, and 60% in water for 1 hour. The samples were decolorized in 75% Methanol / 20% DMSO / 3% Hydrogen peroxide (Sigma Aldrich) at 4°C overnight. Decolorized samples were then washed with 100% Methanol for 1 hour to remove the peroxide and then delipidated in 100% dichloromethane (DCM) twice for 1 hour each, followed by overnight incubation at 4°C in the same solution. The next day, the samples were decalcified with Morse solution (20% formic acid (Carl Roth, Cat no. 5355.1), 10% sodium citrate (Carl Roth, Cat no. 2611.3)) overnight at 4°C. After decalcification, the bone samples (ranging from 3x3x3 mm to 1cm x 0.5cm x 0.5cm in size) were further trimmed to approximately 0.5cm x 3mm x 3mm with RNASE AWAY (Thermofisher, Cat. No. 10328011) cleaned scalpel. The samples were washed with 2x SSCT two times 1 hour each at 4°C. Next, the tissue was permeabilized with permeabilization buffer (0.2% CHAPS (Carl Roth, 1479.4) / 0.1% Deoxycholate, (Carl Roth, Cat no. 3484.1) / 10% DMSO / 25 mM EDTA pH 8.0 in 1x PBS) at 4°C overnight. All pretreatment steps were performed in a scintillation vial (Fisher Scientific, Cat. No. 11526325) with 20 mL of solution with agitation.

##### (ii) 3D RNA detection with Hybridization Chain Reaction (HCR)

Pretreated samples were prehybridized with hybridization buffer (30% formamide, 5x sodium chloride sodium citrate (SSC), 9 mM citric acid (pH6.0), 0.1% Tween 20, 50 µg/mL heparin, 1x Denhardt’s solution, 10% Dextran sulfate, from Molecular Instruments) for 2 hours on a 37°C shaker at 200 rpm (rotations per minute). The samples were hybridized with HCR oligos at 2 pmol per 100 µl of hybridization buffer for 2 days at 37°C. After hybridization, the samples were washed three times, 1 hour each, with wash buffer (30% formamide, 5x sodium chloride sodium citrate (SSC), 9 mM citric acid (pH 6.0), 0.1% Tween 20, 50 µg/mL heparin, from Molecular Instruments), followed by overnight washing with wash buffer on a shaker at 200 rpm at 37°C. Washed samples were incubated with amplification buffer (5x sodium chloride sodium citrate (SSC), 0.1% Tween 20, 10% Dextran sulfate, from Molecular Instruments) at room temperature on a roller mixer for 8 hours, followed by 2 days of HCR amplification at room temperature. The samples were washed with 5x SSCT (5x sodium chloride sodium citrate (SSC), 0.1% Tween 20) three times, 1 hour each, at room temperature, followed by overnight washing with 5x SSCT. Next, the samples were rinsed 2 times in PBS / 0.2% Tween-20 / 10 µg/mL heparin / 25 mM EDTA for at least 1 hour and were ready for subsequent immune-probing or direct clearing. All steps were performed with gentle shaking if not specified otherwise.

The HCR amplifiers and probes were purchased from Molecular Instruments, Inc. (https://www.molecularinstruments.com). Some probes were designed using the HCR probe generator, which can be found here https://github.com/rwnull/insitu_probe_generator, and purchased from IDT. The probe pair sequences can be found in the supplementary information, Supplementary Tables S2 and S3, for reproducibility.

##### (iii) 3D Immunostaining

The tissue was blocked with staining buffer (PBS / 0.2% CHAPS / 5% Donkey Serum (Merck, Cat no. D9663) / 10% DMSO / 25 mM EDTA pH 8) for 8 hours at 37°C. Samples were stained with antibodies in staining buffer for 48 hours at 37°C. Next, the samples were washed in PBS / 0.2% Tween-20 / 10 µg/mL heparin / 25 mM EDTA (HET-PBS) three times, 1 hour each, at room temperature, followed by overnight washing with HET-PBS. The samples were subsequently stained with secondary antibodies in the staining buffer for 48 hours at 37°C. Stained samples were then washed again with HET-PBS three times for 1 hour each, followed by overnight washing in HET-PBS at 37°C. The source of primary and secondary antibodies was documented in Supplementary Table S2. Cell nuclei were stained with YO-PRO™-1 Iodide (Thermofisher, cat. No. Y3603) added during the last overnight washing with secondary antibodies. The bone was visualized based either on eosin staining (excited at 488 nm, acquired with a 525±50 nm emission filter) or autofluorescence signal upon excitation at 405 nm and acquisition with a 460±40 nm emission filter set^31^. As an example, an imaging containing five channels (autofluorescence, YOPRO, CD271, CXCL12, and Endoglin (ENG)) was made by staining CXCL12 mRNA with HCR using B2-AF647 adapter, followed by CD271 (Host: mouse) coupled to Cy3 anti-mouse secondary antibody and ENG (Host: rabbit) coupled to AF790 anti-rabbit antibody staining. The stained samples were further counterstained with YOPRO-1 for labeling cell nuclei in green fluorescence (excitable at 488nm), and the autofluorescent signal from bone matrix was acquired by excitation at 405 nm.

##### (iv) Tissue Clearing

For the clearing, samples were incubated in 20%, 40% Methanol, and 50% tetrahydrofuran (THF, Sigma Aldrich, 360589) for 1 hour each step, followed by 70% THF overnight. The next day, 70% THF was changed to 80%, 90%, and 100% THF three times each for 1 hour at room temperature, followed by 100% THF overnight. Thereafter, THF was washed out with DCM for 15 minutes twice and cleared in dibenzyl ether (DBE, Sigma Aldrich, 108014) for 1 hour at room temperature. Samples were imaged within 48 hours after being placed in DBE. THF (supplemented with chemical stabilizer) was routinely monitored for peroxide level. Risk assessment of experimental procedures and handling of organic solvents were handled, and records were documented according to Institutional health and safety guidelines.

#### Confocal imaging of HCR- and immunofluorescence-stained tissue sections

For two-dimensional imaging, human bone tissue was processed using the same decalcification and delipidation protocol as described for DeepBone clearing. Samples were dehydrated through an ethanol gradient followed by xylene, and embedded in paraffin. Sections were cut at 6 µm thickness and mounted onto Superfrost® Plus Gold Adhesion Slides (Expredia) to minimize detachment during subsequent steps.

HCR staining was performed using the v3 protocol (Molecular Instruments) according to the manufacturer’s instructions. CXCL12 transcripts were detected using probes with a B2 adapter and B2-AlexaFluor647 amplifiers. After hybridization, sections were washed twice in PBS containing 5 mM EDTA and 0.1% Tween-20 to remove residual 5× SSCT salts.

Following HCR, tissue sections were blocked using the same staining buffer as used in DeepBone, then incubated with primary antibodies against LEPR (conjugated to AlexaFluor488; #MAB867, R&D Systems) or Ki67 (conjugated to AlexaFluor546; SolA15, Thermofisher). Nuclei were counterstained with DAPI.

Imaging was performed using a Nikon Yokogawa Spinning Disk Field Scanning Confocal System. Fluorescent signals were acquired using lasers at 405, 488, 546, and 639 nm with a 40× objective, spanning a z-depth of 10 µm at 2 µm intervals. Raw images were saved in ND2 format and converted to .ims format using Imaris Converter (v10.0) for visualization and image export.

#### Lightsheet microscopy and 3D image acquisition

DeepBone-cleared human bone samples were immersed in dibenzyl ether (DBE) and imaged using a light-sheet fluorescence microscope (Ultramicroscope II, LaVision BioTec, Bielefeld, Germany) equipped with an Olympus MVX-10 zoom body (0.63× to 6.3× range) and an Olympus MVPLAPO 2×/0.5 NA objective lens, fitted with a 5.7 mm dipping cap and correction optics. The system provides an axial resolution of approximately 4 μm.

Fluorescence excitation was achieved using Coherent OBIS lasers (488-100 LX, 561-100 LS, and 640-100 LX) and appropriate emission filters. Images were acquired using an Andor Neo sCMOS camera (2560 × 2160 pixels) with a voxel resolution of 0.944 × 0.944 μm in the x-y plane and a z-step interval of 4–5 μm (voxel size: 0.944 × 0.944 × 4 or 5 μm), at a zoom factor of 3.2× (yielding a combined magnification of 6.4×). Image acquisition was performed with ImSpector software (v347), and laser power was adjusted individually to prevent signal saturation. Images were saved as 16-bit TIFF files.

For larger regions, tile scans were performed with a 10% overlap along the longitudinal (y) axis. Tiles were stitched using a non-rigid implementation of Terastitcher (available at: https://github.com/mcrewcow/Chu_et_al_2023/blob/main/terastitcher_ImSpector_.py.). After stitching, the resulting multi-channel z-stacks were converted into .ims format using Imaris Converter (v9.0.1 or v10.0.1), with appropriate metadata (voxel size and channel alignment) included during import.

Multichannel acquisitions were processed by converting each fluorescence channel independently, then combined using the “Add Channel” function in Imaris. The following excitation/emission filter sets were used:

405 nm excitation, 480/40 nm emission for bone autofluorescence

488 nm excitation, 525/50 nm emission for YOPRO-1

561 nm excitation, 595/40 nm emission for Cy3 and Alexa Fluor 546

640 nm excitation, 680/30 nm emission for Cy5 and Alexa Fluor 647

785 nm excitation, 845/55 nm emission for Alexa Fluor 750, 790, and IR800

#### Image processing

Images were processed with Fiji, Terastitcher, and Imaris 9.0.1 (Bitplane, Switzerland). Each acquired z-stack was first corrected for uneven illumination with a batch macro applying enhancement of the local contrast algorithm in Fiji and median filter in the “image processing” option (3x3x3) in Imaris. Movies were prepared using the “animation” function in Imaris and exported in MP4 format. Rates of 24 to 96 frames per second were used.

#### Image processing and vessel segmentation

Image denoising was performed based on published literature to remove 3D volumes that are too large or too small^68^. A cubic volume of approximately 1 mm^3^ was selected away from the edge of the sample (in which the quantification was later normalized by tissue or image frame volume). A surface mask was first generated for each channel based on the “surface” function in Imaris. For the bone surface, only objects with volume under a surface larger than 20,000 µm^3^ were considered to avoid non-specific auto-fluorescent signals. For blood capillaries and sinusoids, noise signals with relative surface sphericity smaller than 0.7 and surface volume larger than 10,000 µm^3^ were considered as previously described^68^ to eliminate subcellular-sized segments. Images of the vessels were further processed with a 3x3x3 median filter function provided by Imaris. Thresholds were optimized for each experimental group to minimize the background in the segmented objects. The numerical values were exported to GraphPad Prism 8.0 for statistical analysis. Images were processed with Windows 10 Pro for Workstations. All data were stored and analyzed using Seagate 4TB External SSDs connected via USB 3.2 or USB 3.2 Type-C ports. Detailed description of the steps can be found in **the Supplementary Methods Files** section “Denoising of Large Artifacts”.

#### 3D cell identification

Distances between cells and structures were analyzed with the “surface” function, “spot” function, and “spatial vantage” option in Imaris v9.0.1 For analysis of individual cells, 3D volumes that are larger than standard cell size were rejected by a volume filter as previously described (>5000 µm^3^)^68^. Each stained cell was analyzed by the “spot” function to generate a class of objects in Imaris. Local (or reverse) cell density was assessed by calculating the average distance to the nine nearest neighbors of the same class of spots.

Total cell density was obtained by first quantitation of cell spots with the “spot” function followed by normalization of the total number of cell spots detected to total tissue volume. Thresholds were optimized for each experimental group with a standardized cell diameter of 12 µm to minimize the background in the segmented objects. Secondary antibody precipitates were minimized by centrifugation of secondary antibodies at 13000 rpm as recommended (https://idisco.info/wp-content/uploads/2015/04/whole-mount-staining-bench-protocol-methanol-dec-2016.pdf). Detailed steps of cell detection can be found in the Supplementary Methods file section “Spot detection pipeline in Imaris.

#### Quantification of Spatial distance from cells to vessels

The distance of spots to the surface was analyzed with MATLAB Xtension package in Imaris. For estimating the percentage cumulative distribution to the surface, the saturation of a particular spot class was analyzed starting from the surface object until all the spots were found within a certain distance from the surface. Colocalization of spots was analyzed with Imaris Xtension packages named “Colocalize Spots”. The distance of a particular cell, identified by spot function mentioned above, to vessel surface distance were calculated by “find spots close to surface” available in the Xtension packages in the spots or surface functional tabs. After these computations. Specific information such as the distances of each spot to vessel surface were exported in the “details: tab within the spot class. For calculation of percentage Cumulative distribution Curves, the statistics were obtained using the “Vantage” option within Imaris 9.0.1. Within the “Spatial View” tab. The tabulated statistics for the percentage saturation of spots(cells) to a particular surface (for example, Eng+ vessels) can be exported as an Excel file. These tabulated files were then input to Prism from independent patients and plotted using “exponential plateau” within “non-linear curve fitting” from biological replicates. Detailed steps of cells-to-surface distance calculation can be found in the Supplementary Methods File.

#### Quantification of CDH13 vessel termini relative to bone and sinusoidal surfaces

To measure the spatial proximity of CDH13+ vessel termini to the bone surface and ENG+ sinusoids in 3D, three regions of interest (ROIs) measuring 400 × 400 × 400 µm were selected from each biopsy sample (∼3 × 3 × 3 mm total volume), avoiding tissue edges and chosen after visual inspection of the CDH13+ vasculature.^22^

Filament objects were generated in Imaris v9.9.1 using the Filament Tracer module. The spatial coordinates of the terminal filament points (classified as “Dendrite Termini”) were exported from the “Position” field in the “Details” tab. These coordinates were then re-imported into the same 3D dataset using the “Import spots from CSV” MATLAB script available from the UCSF BIDC resource (https://bidc.ucsf.edu/create-spots-csv).

To exclude artifacts from ROI cropping, spot objects corresponding to filament termini were filtered to exclude any located within 50 µm of the image borders in x, y, or z dimensions. The resulting terminal points were then analyzed using the same 3D distance quantification workflow described above (see *Quantification of spatial distance from cells to vessels*), yielding minimum distances between CDH13+ vessel termini and either the bone surface or the ENG+ sinusoidal surfaces.

### Calculation of 3D Gaussian Kernel density

For the estimation of the 3D Gaussian Kernel Density, the *x*, *y*, and *z* coordinates of every cell and every point of the vascular/bone surface were exported from Imaris and loaded into R. The following equation was used to calculate the Gaussian kernel density:

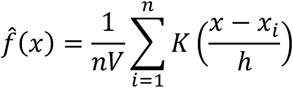

where f^ (x) is the estimated density at point *x*, *n* is the number of data points, *V* is the volume of the 3D space, *x_i_* represents a data point, *K* is the Gaussian kernel function, and *h* is the bandwidth (smoothing parameter). The Gaussian kernel function K was calculated using this formula:

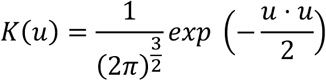

where *u* is the argument vector (*x*/*h*, *y*/*h*, *z*/*h*) in 3D space, and the bandwidth *h* controls the smoothing of the kernel. A smaller *h* results in a sharper density estimate, while a larger *h* produces a smoother estimate. The code for performing density analysis is available at https://github.com/mcrewcow/Chu_et_al_2023/blob/main/3Dcontour.R

### Data preprocessing for Neural Network analysis

The multiplexed 3D images acquired through the lightsheet microscope were converted to spots using the “Spots” function in the Imaris software. Each channel, representing distinct cell type or structure, namely CXCL12, Bone and Endoglin+ sinusoids, was transformed into spot objects with unique spatial coordinates. Spatial coordinates for all cellular and structural components were exported as csv files. All cells within a spherical region around each CXCL12+ cell were counted as a function of the spherical region radius *r*, obtaining the functions *C*_*X*_(*r*), where the subscript X stands for each cell type. Then, these counts as a function of *r* were normalized using the formula:

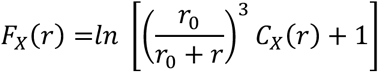

where *r*_0_ is 50 µm is a scaling factor and *r* ranges from 0 to 200 µm in increments of 10 µm. This normalization permits us to control for the volume of the spherical regions where the cells are counted. These functions are then used as inputs for the variational autoencoder (VAE), as described in the next section. The overall dataset is divided into a training set with 90% and a test set with 10% of the data.

### Variational autoencoder architecture (VAE)

The VAE architecture comprises two parts: an encoder and a decoder. For the encoder network, an input layer with 63 neurons (as *r* is a vector of 21 values and there are 3 cell types) leads to a fully connected layer with 16 neurons. This is followed by a Leaky Rectified Linear Unit (ReLU) layer with a 0.2 scalar multiplier for negative input values and another fully connected layer with 4 neurons, culminating in the two-dimensional latent space. The decoder network mirrors the encoder’s architecture, consisting of two fully connected layers and a linear output layer. Sampling the input for the decoder involves the sample layer, which takes as input the mean vector **μ** concatenated with the log-variance vector *ln* (*σ*^2^) and samples elements from *N*(*μ*, *σ*^2^). The log-variance is used to enhance the numerical stability during training.

### ELBO loss function

The Evidence Lower Bound (ELBO) loss function, used for training the VAE, consists of two terms: the reconstruction loss and the Kullback–Leibler (KL) loss. The reconstruction loss *L*_*R*_, measured using the mean-squared error (MSE), evaluates the proximity of the decoder output *x̂*_*i*_ to the original input *x*_*i*_, where *N* is the number of input variables:

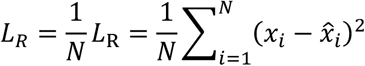

The KL loss *L*_*KL*_, a Kullback-Leibler divergence term, assesses the disparity between two probability distributions. It is calculated as:

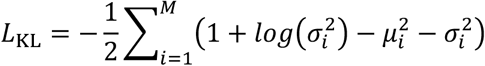

Where *M* is the number of latent variables. Including the KL loss term ensures that clusters, learned through the reconstruction loss, tightly populate the center of the latent space, forming a continuous space for sampling.

The VAE model was trained with a batch size of 128, over 120 epochs, and a learning rate of 10^-3^.

Spatial coordinates of analyzed cells in their corresponding environment in the latent space were retrieved and compiled as csv files. The csv files were imported back to the original 3D image for visualization of cells in their environments, using matlab interface createSpotsFromCSV.m in Imaris from the following source. https://github.com/BIDCatUCSF/Create-Spots-From-Text.

#### Mouse samples

Mouse bones were harvested from 1.5-month-old C57/Black mice. Samples were passively fixed in 4% PFA/1X PBS overnight at 4°C, followed by RNA staining employing protocols and their modifications as indicated above. Male and Female mice were pooled together for analysis. The bone harvesting was ethically approved by the Northern Stockholm Animal Research Committee (ethical permit number: 18160-2019). All animal manipulations were performed according to The Swedish Animal Agency’s Provisions and Guidelines.

#### Analysis of Single-Cell RNA Sequencing Data from Human Bone Marrow

To construct a comprehensive human bone marrow atlas, we merged previously published single-cell RNA sequencing (scRNA-seq) datasets, resulting in a combined dataset of approximately 145,000 cells. Following quality control filtering based on standard cell metrics, ∼80,000 high-quality cells were retained for downstream analysis.

Raw gene expression counts were normalized by sequencing depth per cell and log1p-transformed. We identified 3,000 highly variable genes. Reads that were subsequently scaled and clipped before performing principal component analysis (PCA) for dimensionality reduction. Batch effects across datasets were corrected using the Harmony algorithm^69^, applied at the level of principal components. The first 25 principal components were used to construct a neighborhood graph, with each cell connected to its 50 nearest neighbors.

Unsupervised clustering was performed using the Leiden algorithm, yielding distinct clusters that broadly corresponded to major cell types. For visualization, we employed Uniform Manifold Approximation and Projection (UMAP). Differential gene expression analysis was conducted using the Wilcoxon rank-sum test to identify genes significantly upregulated in each cluster compared to all other cells, with multiple testing correction applied via the Benjamini-Hochberg method.

Stromal subclustering: Stromal cluster, together with the adjacent osteo-lineage cluster, was isolated for further analysis. To resolve finer subtypes within each population, we applied Leiden clustering independently to each cell type. Characteristic genes for the resulting subclusters were identified using the same Wilcoxon-based approach.

Endothelial cells: To investigate RNA expression across different vascular subtypes, we compiled an additional dataset by selecting a well-defined endothelial cluster (PECAM1+, KDR+, EMCN+) from our atlas, supplemented with endothelial cells from two publicly available human bone scRNA-seq datasets^13,70^. This yielded a total of approximately 6,000 endothelial cells.

Due to the complexity of primary human transcriptomic data, we utilized the scVI module for data processing and integration. To further enhance cross-species comparison, we employed the MultiMap integration tool to align the human dataset with previously published and well-annotated murine bone marrow vessel transcriptomes^54^. Detailed methodology is provided in the Supplementary Methods File. All scRNA-seq analyses were conducted within the Python-based scverse ecosystem.

#### Statistical analyses

Unpaired Student’s t-test was applied everywhere where only two groups were compared from different patients. When groups from the same set of patients were compared, a paired t-test was employed. For non-normally distributed data, the Mann-Whitney U test was employed to compare independent datasets, and the Wilcoxon matched-pair signed rank test was employed to compare linked datasets. To compare multiple groups, one-way ANOVA with multiple comparisons was performed. Statistical tests for the percentage cumulative distribution of cells to surface were performed using the Kolmogorov-Smirnov test. All statistical analyses were performed with at least three biological (human samples) replicates. For all statistical tests, a minimally acceptable significance level at p<0.05 was considered. No statistical method was used to predetermine sample group sizes. Values are presented as mean or median with Standard Error for error bars. No randomization or blinding was used for animal studies. Human samples were randomly coded to avoid bias.

## Reporting Summary

Further information on research design is available in the Nature Research Reporting Summary linked to this article.

## Data Availability

The human bone marrow atlas can be explored at the CellxGene Portal: https://cellxgene.cziscience.com/collections/0391c84c-d57d-4741-9277-e4d58f9a3d0c

The main data supporting the results in the study are available in supplementary information, some of the key scans are open to public access via the following link https://zenodo.org/uploads/14212791?token=eyJhbGciOiJIUzUxMiJ9.eyJpZCI6IjUyNTgxZTc3LTI1ZWItNDk3ZC1hN2M4LWJkMThlOGFhZTRlMCIsImRhdGEiOnt9LCJyYW5kb20iOiI4MGY2NzY4MGFmNjM1YmYyYTA3OTQxMzA5NDViMzc1OCJ9.i_P3vGu03LP8-Yg1VdArRqzt1x6fVzE0_l9iQxH2leVH3_8cDQkid4rXwyI4aD_Oj34ggaYrvTfjFVSq6I95fg descriptions of the files are included in the above repository.

Other raw and analysed datasets generated during the study are too large to be publicly shared, yet they are available for research purposes from the corresponding authors on reasonable request.

## Code Availability

The code generated in this study is deposited to GitHub and available by the following link: https://github.com/mcrewcow/Chu_et_al_2023

## Acknowledgments

We thank patients and their parents for donating their tissue for research purposes. We thank Olga Kharchenko for the illustrations. The data handling was enabled by resources in project NAISS sens2023024 provided by the National Academic Infrastructure for Supercomputing in Sweden (NAISS) at UPPMAX, funded by the Swedish Research Council through grant agreement no. 2022-06725. The computations were enabled by the Berzelius resource in project Berzelius-2023-108 provided by the Knut and Alice Wallenberg Foundation at the National Supercomputer Centre.

## Author contributions

A.S.C. and N.C.T. conceived the study. N.C.T performed the majority of the experiments with the help of E.K., D.T., M.S.Z., L.L., Z.L., X.L., X.T.. O.D. carried out all the bioinformatics analysis. I.G., C. O., M.B., J.B.P., R.H.A., and G.V. contributed intellectually throughout the project. L.S., I.G., M.B., and B.B. provided the human samples. Y.W.C., E.C.R., and G.V. provided the neural network analysis pipeline. N.C.T. and A.S.C. wrote the manuscript. All of the authors critically reviewed this manuscript and approved the final draft.

## Competing interests

The authors declare no competing interests.

**Video S1: Visualization and quantification of the vasculature in the aged human bone marrow.** Aged human subchondral bone (aged 74 years old) was processed with *DeepBone,* and vasculature was stained using CD31 antibody. Vessels were computed with the Imaris “filament” function, and individual filaments were color-coded by filament diameter from blue (thin) to red (large).

**Video S2: *DeepBone* enables the detection of mRNA transcripts inside the human bone matrix** Detection of SOST mRNA (red) throughout an aged human bone with *DeepBone*. Cyan represents autofluorescence at 405 nM.

**Video S3: Spatial arrangement of BM-MSCs, proliferating cells, and capillaries in young human bone.**

Adolescent human subchondral bone (14 years old) was processed with *DeepBone,* and was stained for CXCL12 mRNA (BM-MSCs, red), CD31(blood capillaries, yellow), and Ki67 (proliferating cells, white).

**Video S4: *Spatial arrangement of BM-MSCs, capillaries, and adipocytes in young human bone*.** Human bone from an adolescent individual (14 years of age) was stained for CXCL12 mRNA (BM-MSCs, red), CD31 (blood capillaries, yellow), and perilipin (adipocytes, white).

**Video S5: *Spatial arrangement of BM-MSCs, capillaries, and adipocytes in aged human bone*.** Human bone from an aged individual (74 years of age) was stained for CXCL12 mRNA (BM-MSCs, red), CD31(blood capillaries, yellow), and perilipin (adipocytes, white).

**Video S6: *2D optical sectioning of BM-MSCs, capillaries, and adipocytes in entire aged human bone biopsies.***

Human bone from an aged individual (74 years of age) was stained for CXCL12 mRNA (BM-MSCs, red), CD31(blood capillaries, yellow), and perilipin (adipocytes, white).

**Video S7: Gland-like appearance of adipocytes in adolescent human bone.** Human bone from an adolescent individual (14 years of age) was stained as in Video S3 but presented with a higher magnification focusing on the adipocyte-rich region White – adipocytes, yellow – blood vessels, red – BM-MSCs, green - YOPRO-1 counterstain.

**Video S8: The canopy membrane in aged human bone.** The canopy membrane was visualized in aged human bone (74 years of age) utilizing immuno-detection of CD271 (yellow). Sinusoids were visualized by Endoglin antibodies (white) and bone by autofluorescence at 405 nM (cyan).

**Video S9: Sinusoidal vessels lining the bone surface in young human bone, making a special niche for BM-MSCs.**

Sinusoids were immune-detected by Endoglin antibodies (white), BM-MSCs by CXCL12 mRNA expression (red), and bone matrix by autofluorescence at 405 nM (cyan) in adolescent (∼14 years old) human bone. Purple spots indicate CXCL12**+** cells located within 20µm distance to both bone and endoglin surface.

**Video S10: Osteo-sinusoidal niche for BM-MSCs in young human bones.** Sinusoids were immune-detected by Endoglin antibodies (white), BM-MSCs by CXCL12 mRNA expression (red), and bone matrix by autofluorescence at 405 nM (cyan) in adolescent (∼14 years old) human bone. Purple spots indicate CXCL12**+** cells located within 20µm distance to both bone and endoglin surface.

**Video S11: Osteo-sinusoidal niche for BM-MSCs in aged human bones.** Sinusoids were immune-detected by Endoglin antibodies (white), BM-MSCs by CXCL12 mRNA expression (red), and bone matrix by autofluorescence at 405 nM (cyan) in adolescent (∼14 years old)

**Extended Data Figure 1.**
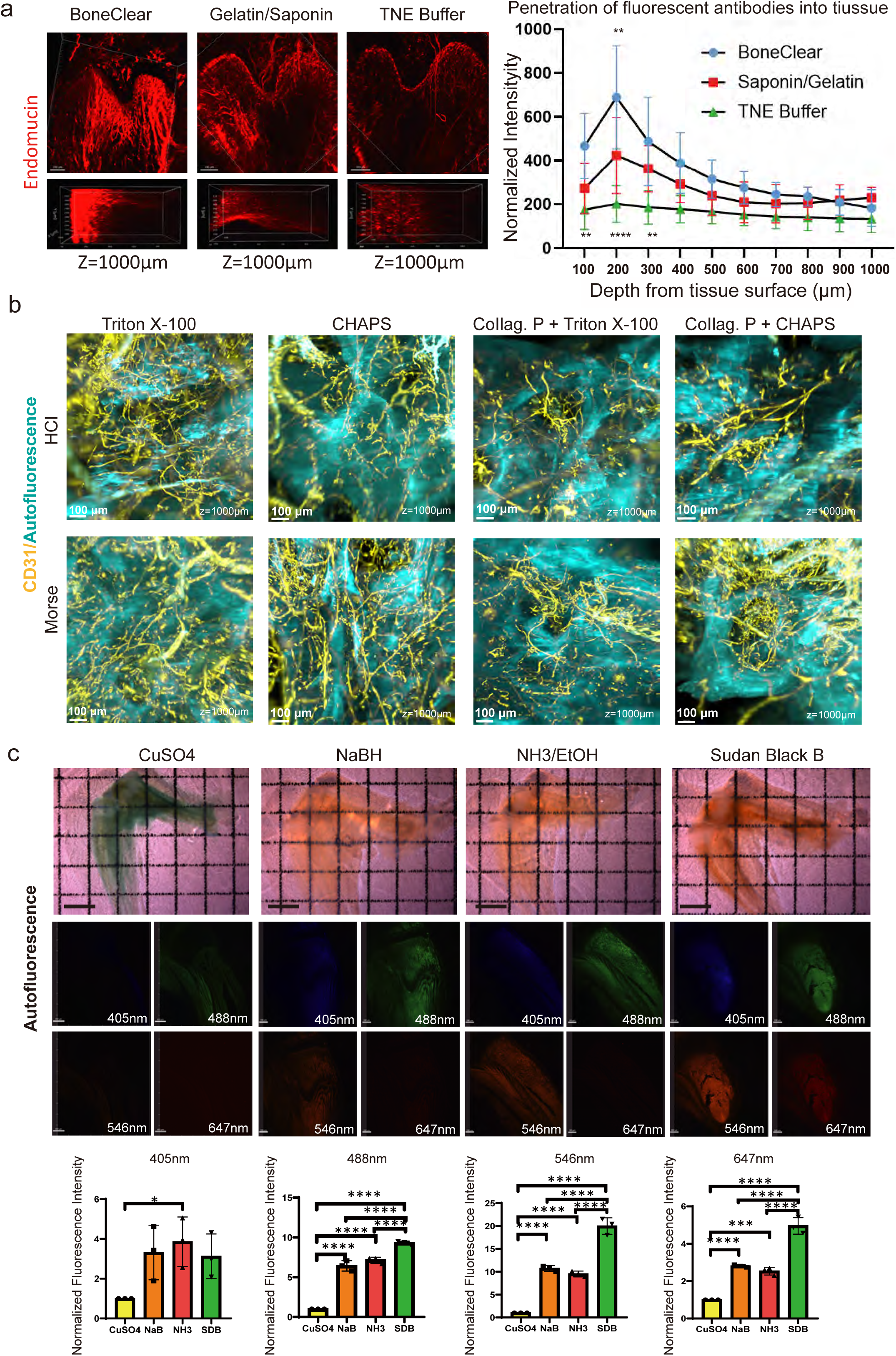
Optimizing protein detection in bone samples. **a**, Representative immunostaining of endomucin, a marker of the sinusoidal vasculature in the murine tibia, using previously published protocols, along with a graph quantifying the depth of the fluorescence achieved. All bones were scanned up to 1 mm in depth (z-axis). The statistical analysis was performed by one-way ANOVA, and mean±SD values are shown on the right panel, n=3. **b**, Testing of different decalcification and permeabilization approaches on aged human bone combined with immunofluorescent detection of CD31 (CD31 – yellow, autofluorescence – blue). Collag. = collagenase P. **c**, Comparison of the ability of CuSO4, sodium borohydride (NaBH), ammonium (NH3 in 70% ethanol), and Sudan Black B (SDB) to inhibit the autofluorescence of bone matrix. Autofluorescence was detected with a Lavision microscope following excitation (at equal power) with light at 405, 488, 546, and 647 nm. Representative images are shown, along with quantification (mean±SD, n=3) of the autofluorescence obtained under various conditions. Statistical analysis was performed by one-way ANOVA.

**Extended Data Figure 2.**
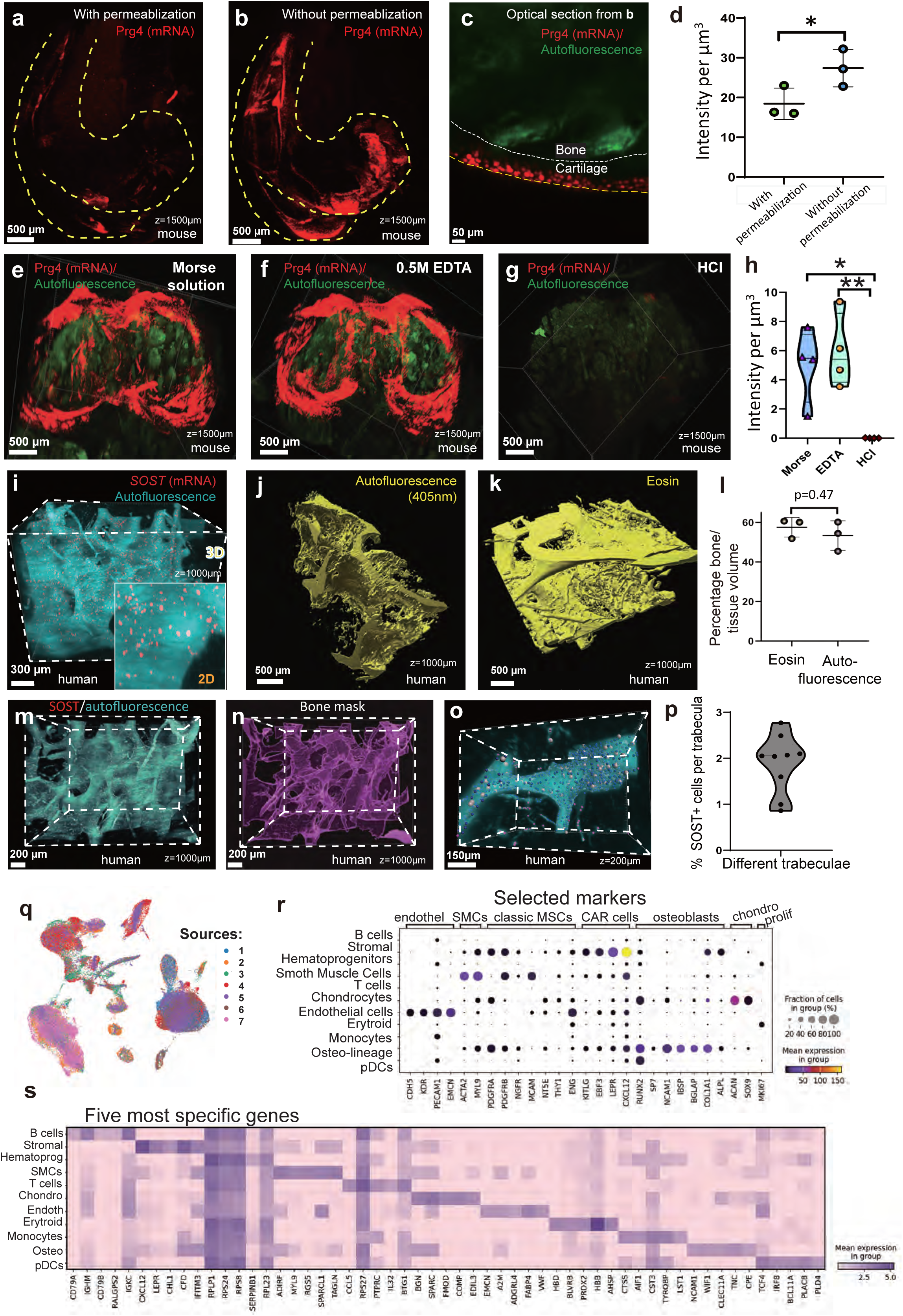
Optimization of mRNA detection in bone tissue and integration of scRNA-seq datasets. **a–c**, Detection of Prg4 mRNA (red) using HCR RNA-FISH in whole-mounted mouse distal femurs (lateral view) with (**a**) and without (**b**) permeabilization; **c**, corresponding optical section from (**b**). **d**, Quantification of Prg4 signal intensity with and without permeabilization (mean ± s.d., n = 3). Statistical comparison: unpaired t-test; P < 0.05. **e–g**, Effect of 24-hour decalcification using Morse solution (**e**), EDTA (**f**), or HCl (**g**) on Prg4 detection (red) in whole-mounted mouse distal femurs (frontal view). Green signal denotes autofluorescence. **h**, Quantification of Prg4 signal following different decalcification protocols. Statistical comparison: One-way ANOVA; *P < 0.05, ** P < 0.01. **i**, Detection of SOST mRNA (red) expressed by osteocytes embedded within the bone matrix. Cyan: autofluorescence at 405 nm excitation. Inset: optical section. **j–k**, Visualization of bone volume using autofluorescence at 405 nm (**j**) or eosin counterstaining (**k**) on the same sample. **l**, Quantitative comparison of segmented volumes from (**j**) and (**k**); paired two-tailed t-test. **m–n**, Rendering of bone surface based on autofluorescence excited at 405 nm. **o**, Example of 3D segmentation of an individual trabecula and SOST-positive osteocytes detected with Imaris. Cells are colour-coded by distance to bone surface (red: closest; blue: farthest), and size-coded by depth relative to image front. **p**, Percentage of SOST-positive osteocytes within individual trabeculae, normalized to total nucleated cells. **q**, UMAP plot of bone marrow cells from this study merged with six publicly available scRNA-seq datasets: (1) Zhang et al. (ref. 32), (2) this study, (3) Qiu et al. (ref. 33), (4) Li et al. (ref. 12), (5) Liu et al. (ref. 34), (6) Kfoury et al. (ref. 35), (7) Hu et al. (ref. 36). **r**, Bubble plot showing expression of selected markers across non-hematopoietic clusters (cluster identities shown above). Note, normalization is done to the depth of sequencing only. **s**, Heatmap of the top five differentially expressed genes per cluster. Cluster annotations: B cells – B lymphocytes; Stromal – bone marrow mesenchymal stromal cells (BM-MSCs); Hematoprog – hematopoietic progenitors; SMC – vascular smooth muscle cells; T cells – T lymphocytes; Chondro – chondrocytes; Endoth – endothelial cells; Erythroid – erythroid lineage; Osteo – osteolineage; pDC – plasmacytoid dendritic cells.

**Extended Data Figure 3.**
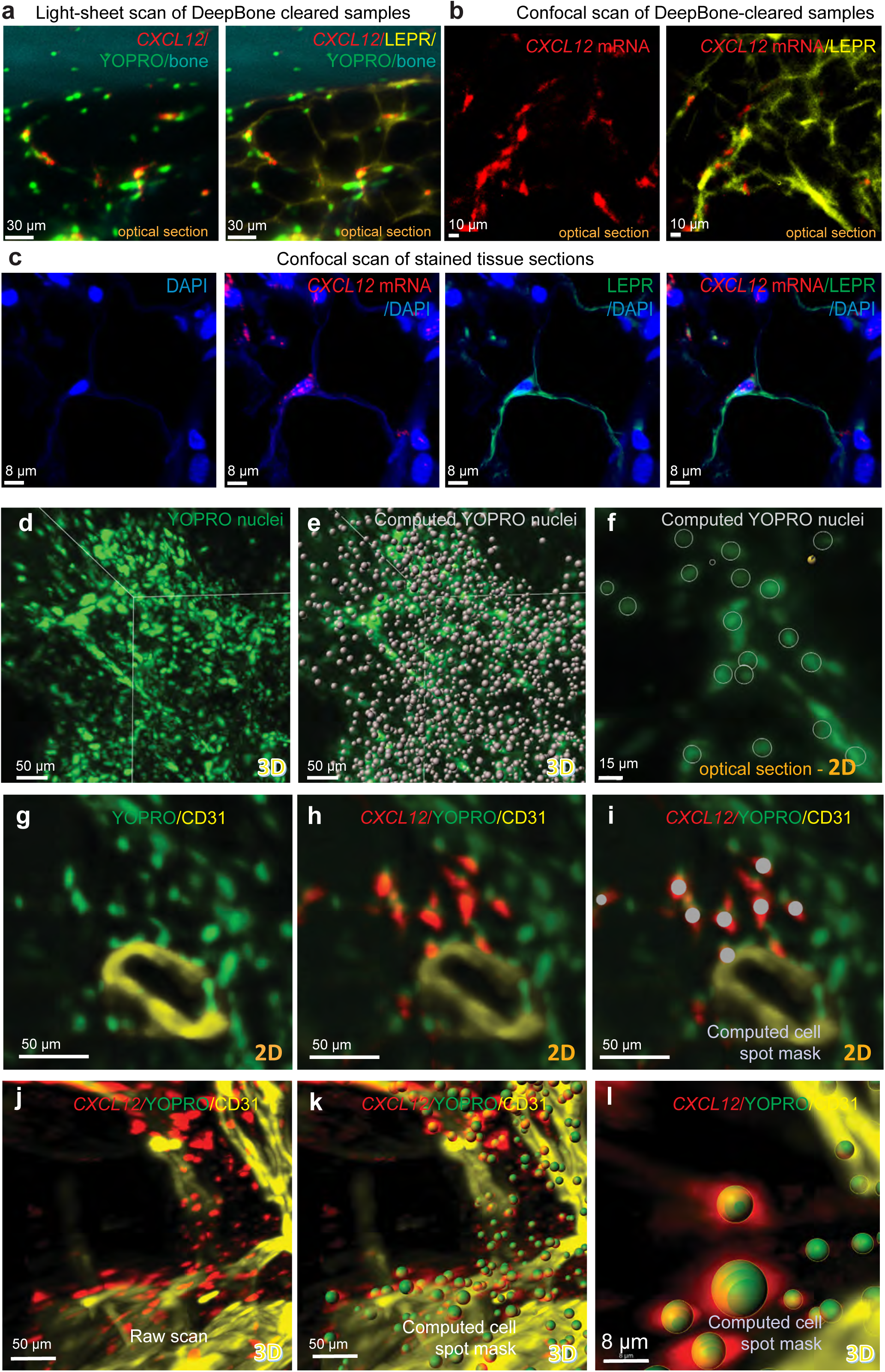
Validation of stromal cell identity and detection strategy. **a,b**, Colocalization of LEPR protein (antibody staining, pseudocoloured yellow) and CXCL12 mRNA (HCR RNA-FISH, red) in representative 2D optical sections from (**a**) a 3D light-sheet scan and (**b**) a confocal scan of tissue-cleared human bone. Left panels show LEPR omitted; right panels include all channels. Nuclei (YOPRO, green) and bone matrix (autofluorescence at 405 nm, cyan) were visualized in (**a**) but not (**b**) due to sample bleaching. **c**, Confocal image of a 6-μm thick section from decalcified human bone stained for LEPR (green), CXCL12 mRNA (red), and counterstained with DAPI (blue). Single and merged channels are shown. **d–f**, Nuclear staining of aged human bone with YOPRO1 (green). d, Raw YOPRO+ signals in a light-sheet scan. e, YOPRO+ nuclei overlaid with cell positions detected by the Imaris Spot function. f, Optical sections showing overlay of YOPRO+ nuclei with segmented cell masks (grey circles). **g–l**, Detection of nucleated CXCL12+ cells in aged bone. **g–i,** Representative 2D optical sections from a 3D light-sheet scan showing CXCL12 mRNA (HCR, red), CD31+ vessels (yellow), and YOPRO1+ nuclei (green). CXCL12 signal is omitted in **g** and present in **h**; **i**, colocalized CXCL12+YOPRO+ cells detected by Imaris Spot function are marked by grey circles. **j–l**, 3D projections of the same region as a raw scan (**j**) and detected double-positive cells at low (**k**) and high (**l**) magnification, with double-positive cells represented as green spheres. **j**, CD31 and YOPRO channels only; **k,l**, all channels including CXCL12. All images are cropped from large tissue volumes (∼2 × 2 × 2 mm) scanned by light-sheet microscopy.

**Extended Data Figure 4.**
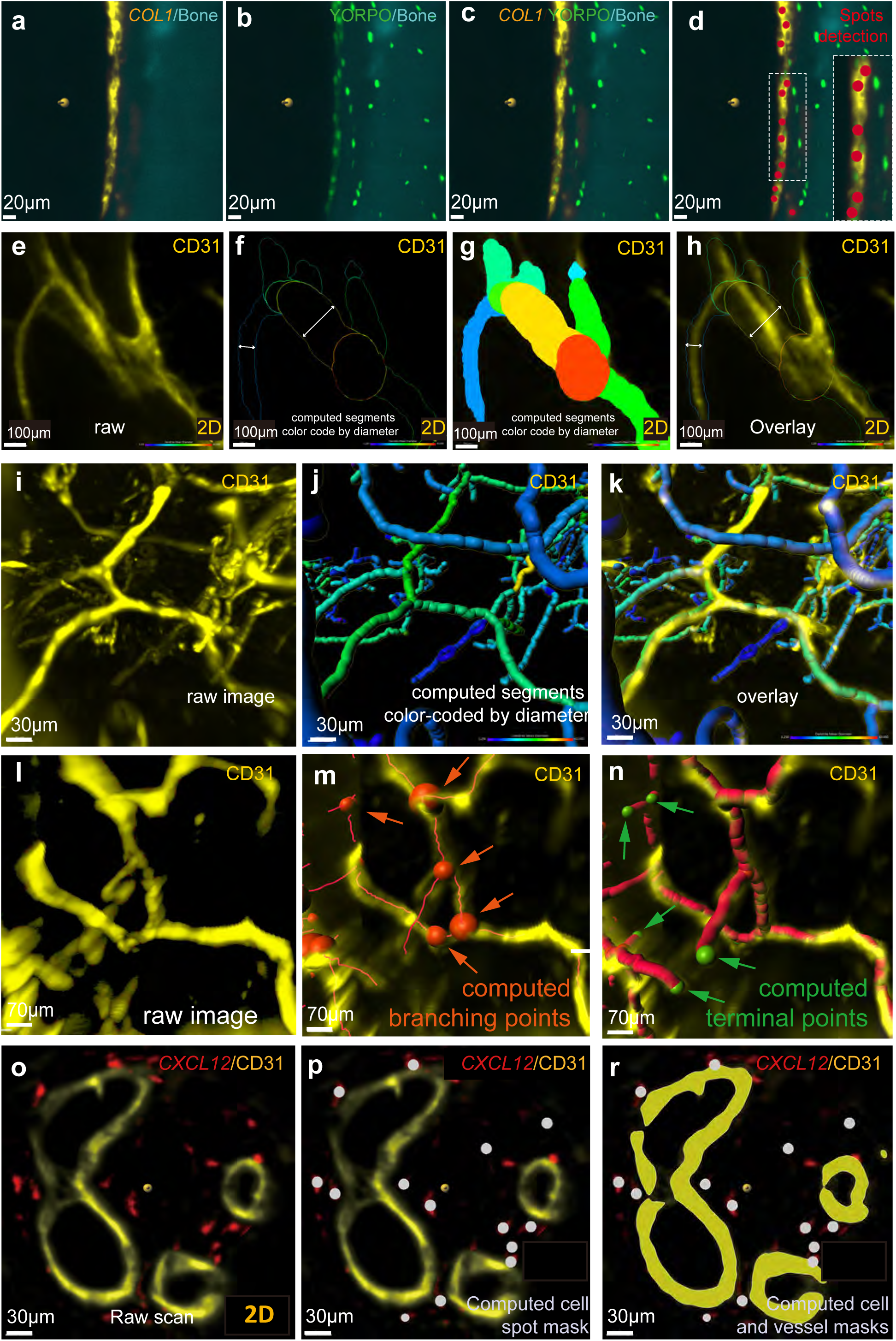
Quantification of tightly packed cells and vascular segmentation in cleared human bone. **a–d**, Detection and segmentation of COL1A1+ cells in aged human bone. **a–c**, Representative optical sections from a light-sheet scan showing COL1A1 mRNA (HCR, yellow), YOPRO1-labeled nuclei (green), and bone matrix autofluorescence (cyan, 405 nm excitation). **d**, Overlay of computed spot mask (red circles) with raw signal, showing automated detection of COL1A1+ cells obtained upon threshold optimization and visual inspection. **e–n**, Segmentation of CD31+ vasculature using filament tracing in Imaris. **e–h**, Optical sections showing raw CD31 signal (**e**), thresholded segmentation (**f**), 2D filament mask colour-coded by vessel diameter (**g**), and overlay of raw and segmented vessels (**h**). **i–k**, 3D projections of a selected volume showing raw CD31 signal (**i**), diameter-coded vessel filaments (**j**), and overlay (**k**). **l–n**, Identification of branching and terminal points within segmented vessels. **l**, Original CD31 signal; **m**, segmented filaments (red lines) and branch points (red spheres); **n**, terminal points (green) and segmented vessels (red) overlaid on raw CD31 signal. **o–r**, Combined spot detection and vessel segmentation. **o**, Optical section showing CXCL12 mRNA (red) and CD31 (yellow); **p**, spot detection of CXCL12+ cells (grey circles); **r**, overlay of cell mask and vascular surface presented using the Imaris Surface function applied on the segmented CD31+ vessels.

**Extended Data Figure 5.**
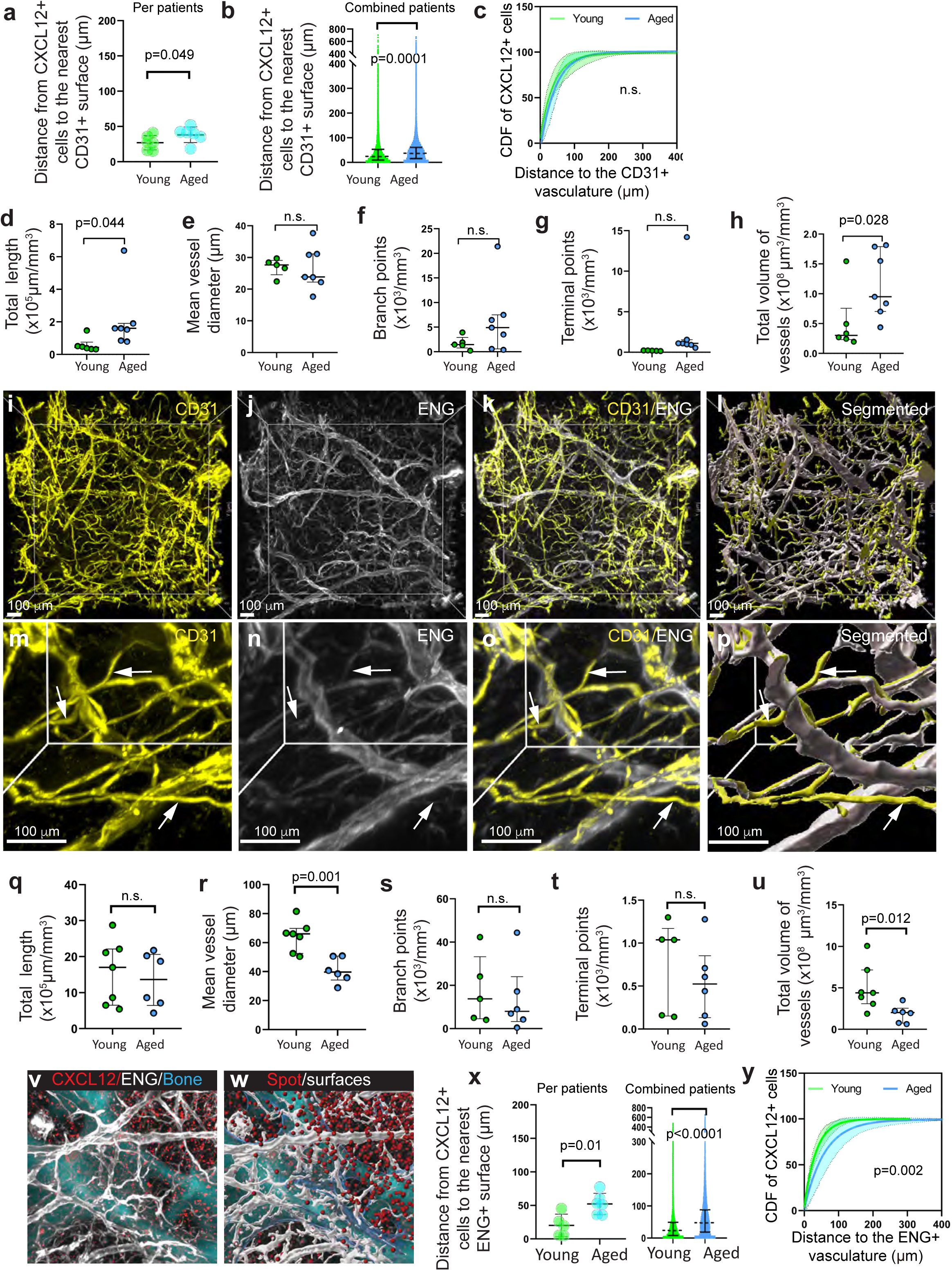
Quantitative spatial analysis of CD31+ and ENG+ vasculature and proximity of CAR cells in young and aged human bone. **a–c**, Distance measurements between CXCL12+ cells and CD31+ vasculature in biopsies from young (green, n = 6) and aged (blue, n = 6) individuals. **a**, Mean per patient (n=patient); **b**, violin plots showing the distribution of all quantified cells (n=cells); dashed lines indicate medians. Statistical comparisons: unpaired t-test (**a**) and Mann–Whitney U test (**b**). **c**, Cumulative distribution functions (CDFs) of distances between CXCL12+ cells and CD31+ capillaries in young (green) and aged (blue) bone samples, with 95% confidence intervals shown as shaded areas. Kolmogorov–Smirnov test (n=patients); n.s., not significant. **d–h**, Quantitative analysis of CD31+ vasculature in young (n = 5) and aged (n = 6) bone biopsies: total length (**d**), mean diameter (**e**), number of branch points (**f**), number of termini (**g**), and total vascular volume (**h**). Data in **d, f–h** are normalized to tissue volume. Statistical analysis: unpaired t-test. **i–p**, 3D co-visualization of CD31+ (yellow) and endoglin (ENG)+ (white) vasculature in aged bone. Low (**i–l**) and high (**m–p**) magnification views are shown. Individual channels (**i, m**), merged projections (**k, n**), and surface masks of segmented vessels (**l, p**) are displayed. White arrows highlight thin CD31-bright, ENG-negative capillaries. **q–u**, Quantitative parameters of ENG+ vasculature in young (n = 5–7) and aged (n = 6) individuals: total length (**q**), mean diameter (**r**), number of branch points (**s**), number of termini (**t**), and total volume (**u**). Data in **q, s–u** normalized to tissue volume. Unpaired t-test. **v–w**, 3D light-sheet imaging of CXCL12+ cells (red), ENG+ vasculature (white), and bone matrix (cyan, 405 nm autofluorescence). **v**, Raw projection; **w**, overlaid with computed cell spots and vascular surface masks. **x**, Distance of CXCL12+ cells to nearest ENG+ vasculature in young (green, n = 5) and aged (blue, n = 6) bone: per-patient means (left) and pooled violin plots (right); dashed lines mark medians. Statistical comparisons: unpaired t-test (left, n=patients) and Mann–Whitney U test (right, n=cells). **y**, Cumulative distribution functions (CDFs) of CXCL12+ cells relative to ENG+ vessels in young (green) and aged (blue) groups with 95% confidence intervals (shaded). Kolmogorov–Smirnov test (n=patients). The number of patients varies between analyses because of technical issues, and none of the samples were excluded by purpose.

**Extended Data Figure 6.**
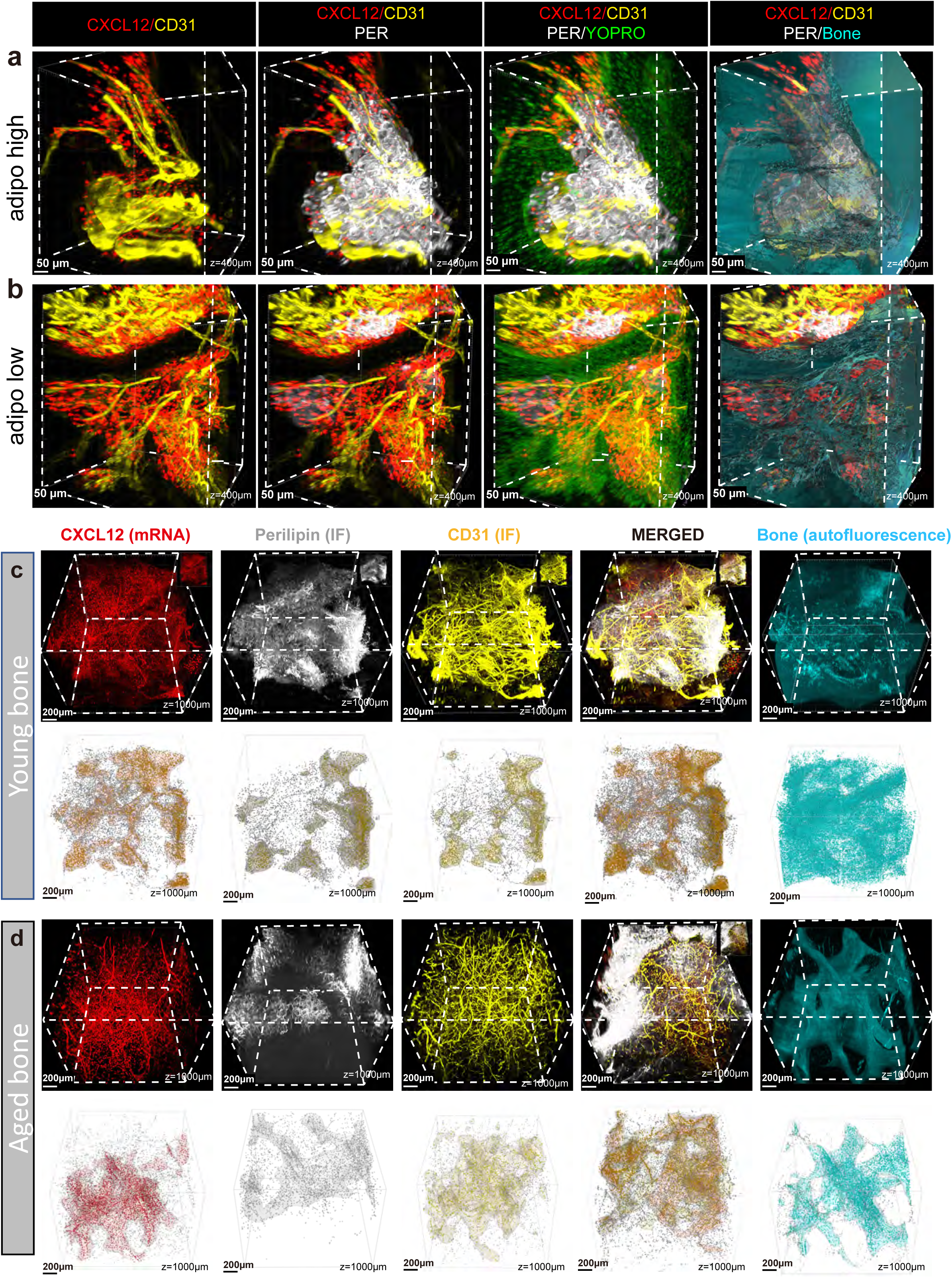
Heterogeneity of human bone marrow. **a**,**b**, Representative images of regions in young bone with high (**a**) and low (**b**) contents of adipose tissue stained for perilipin protein (white, adipocytes), CXCL12 mRNA (red, stromal cells), CD31 protein (yellow, blood vessels), and nucleI (YOPRO nuclear dye, green). **c**,**d**, Three-dimensional Gaussian kernel density map of CXCL12, perilipin, CD31, and bone matrix (autofluorescence) in young and aged human bone.

**Extended Data Figure 7.**
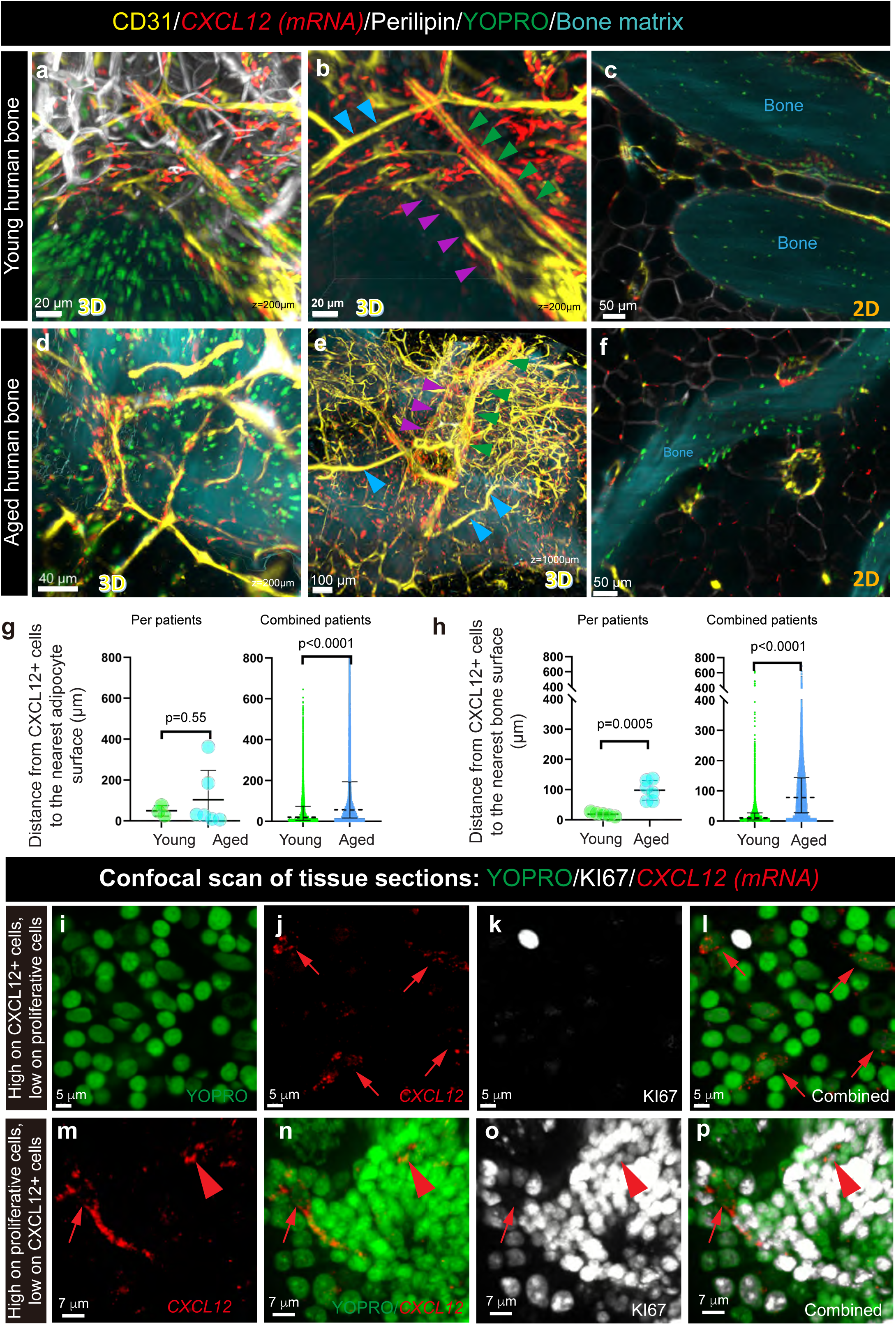
Spatial proximity of CAR cells to adipocytes and bone, and confocal validation of proliferating stromal cells in human bone marrow. **a–f**, Representative 3D projections (**a, b, d, e**) and 2D optical sections (**c, f**) of light-sheet scans of young (**a–c**) and aged (**d–f**) human bone stained for CXCL12 mRNA (red), CD31+ vasculature (yellow), and perilipin+ adipocytes (white). Nuclei were counterstained with YOPRO-1 (green), and the bone matrix was visualized by autofluorescence at 405 nm (cyan). Green arrows: arteries; blue arrows: CD31+ capillaries; purple arrows: CD31-low sinusoids. **g,h**, Spatial distances of CXCL12+ cells to (**g**) perilipin+ adipocytes in young (n = 3) and aged (n = 6) bone biopsies and to (**h**) the nearest bone surface (n = 6 per group). In **g**, **h**, left panels show per-patient means; right panels show violin plots of pooled data. Dashed lines in violin plots indicate medians. Statistical tests: unpaired t-test (left panels, n=patients) and Mann–Whitney U test (right panels, n=cells). **i–p**, Confocal imaging of tissue sections of aged human bones showing marrow regions with low (**i–l**) and high (**m–p**) proliferative activity, stained for YOPRO-1+ nuclei (green, **i, m**), KI67+ proliferating cells (white, **j, n**), and CXCL12 mRNA (red, **k**, **o**). Merged channels are shown in **l** and **p**. Red arrows highlight CXCL12+ cells, and the red arrowhead highlights a double KI67+CXCL12+ cell.

**Extended Data Figure 8.**
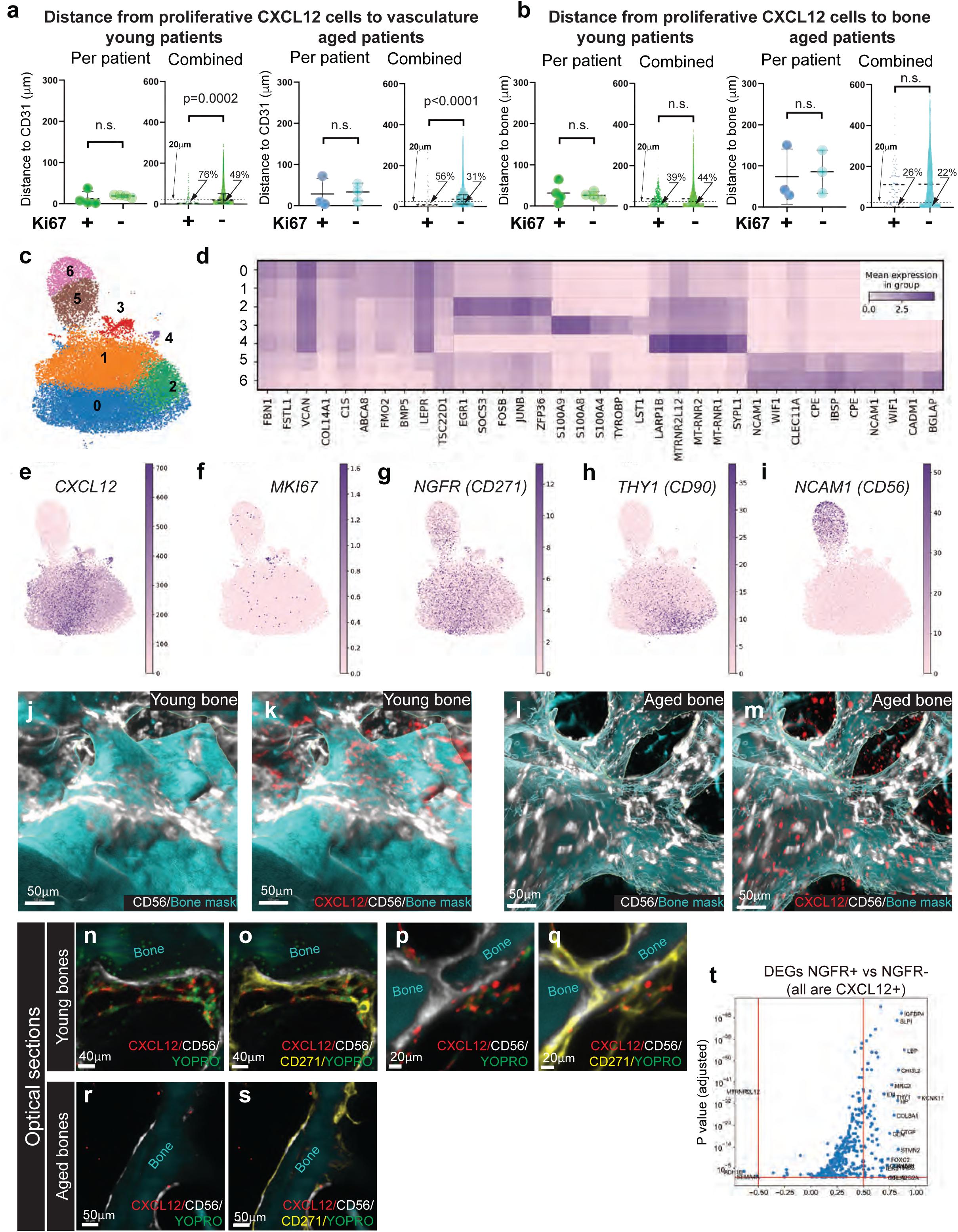
Spatial localization of proliferating CAR cells and detection of osteo-primed CAR subpopulations. **a, b**, Spatial proximity of KI67+ versus KI67– CXCL12+ stromal cells (CAR cells) to CD31+ vasculature (**a**) and bone surfaces (**b**) in young (green) and aged (blue) human bone. Left panels: patient-level means (each dot represents one patient). Right panels: pooled single-cell data displayed as violin plots (dashed lines indicate 20 µm; percentages show the proportion of cells within that threshold). Statistical tests: paired t-test (left, n=patients), Mann–Whitney U test (right, n=cells), n.s. – not significant. **c**, UMAP embedding of stromal and osteolineage clusters (identical to Main Fig. 5a; shown here for orientation). **d**, Heatmap showing the top five differentially expressed genes per cluster from panel **c**. **e–i**, Feature plots displaying normalized gene expression of CXCL12 (**e**), KI67 (**f**), NGFR (CD271) (**g**), THY1 (CD90) (**h**), and NCAM1 (CD56) (**i**) across subclusters. **j–m**, 3D visualization of CXCL12+ (red) and NCAM1+ (white) cells in young (**j, k**) and aged (**l, m**) human bone; bone surface rendered in cyan. CXCL12 channel omitted in **j** and **l** to aid visualization of localization. **n–s**, Representative optical sections from 3D light-sheet scans showing the spatial arrangement of CD271+ (yellow), CXCL12+ (red), and NCAM1+ (white) cells in young (**n–p**) and aged (**r–s**) bone. YOPRO-1+ nuclei (green) and bone autofluorescence at 405 nm (cyan) are also shown. CD271 channel omitted in **n**, **p**, and **r** to aid visualization of co-localization. **t**, Volcano plot showing differentially expressed genes between CD271+ and CD271– stromal populations (scRNA-seq analysis).

**Extended Data Figure 9.**
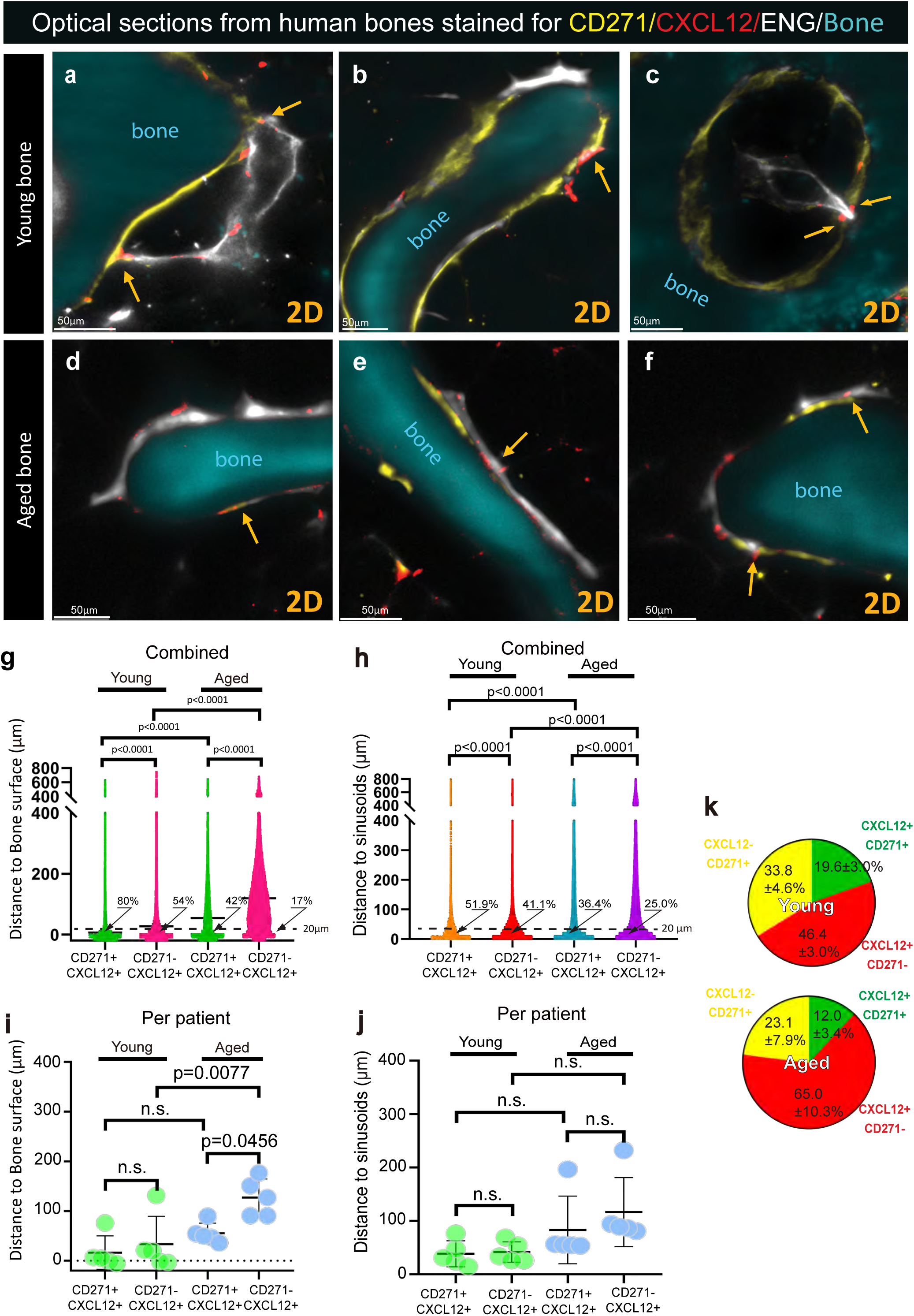
Properties of the CD271⁺CXCL12⁺ stromal subpopulation in human bone. **a–f**, Representative 2D optical cross-sections of sinusoid–bone interactions in young (**a–c**) and aged (**d–f**) human bone biopsies. Immunofluorescence staining of CD271 (yellow) and endoglin (ENG; white), HCR detection of CXCL12 mRNA (red), and bone matrix visualized by autofluorescence following 405 nm excitation (cyan). Orange arrows indicate CXCL12+ cells in proximity to both sinusoids and bone surfaces. **g-j**, Violin plots showing the distance of CD271+CXCL12+ versus CD271-CXCL12+ cells to the nearest bone surface (**g, i**) or sinusoidal surface (**h,j**) in young and aged bone and per-patient means in (**i)** and (**j),** respectively. Dashed lines mark the 20 µm threshold, and percentages denote the fraction of cells within this range. Statistical comparison: one-way ANOVA with n=cells in **g,h**, and n=patients in **i,j**. **k**, Pie charts summarizing the relative proportions of CD27+, CXCL12+, and double CD27+CXCL12+ cell populations, across bone biopsies from young and aged individuals (mean±SE, n=5 patients/age group).

**Extended Data Figure 10.**
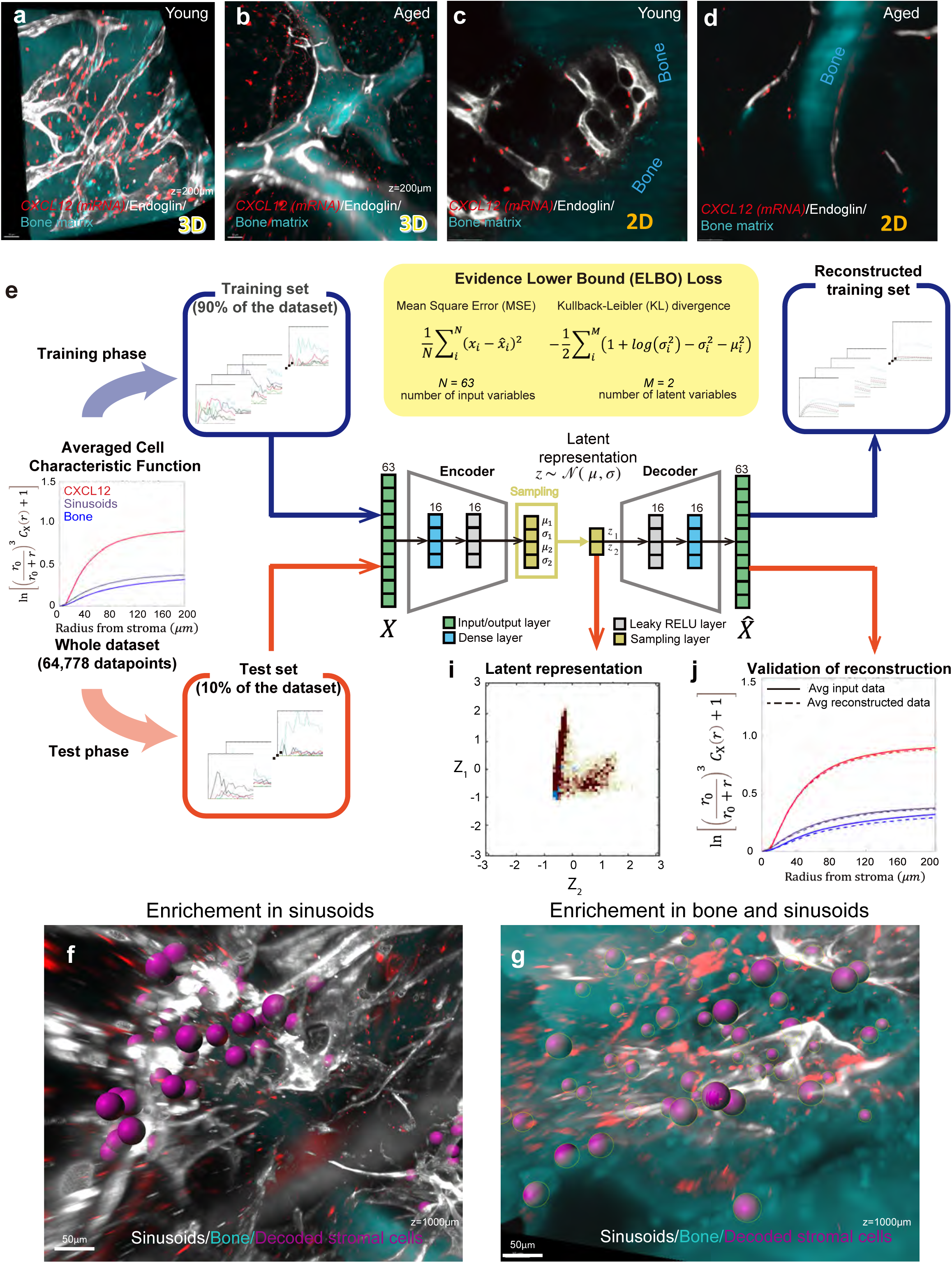
Sinusoidal–bone composites, architecture of the variational autoencoder, and niche reconstruction. **a–d**, Visualization of sinusoidal–bone composites and associated CXCL12+ cells in young (**a, c**) and aged (**b, d**) human bone samples. Sinusoids (endoglin, ENG; white), CAR cells (CXCL12 mRNA, red), and bone matrix (autofluorescence at 405 nm excitation, cyan) are shown in 3D projections (**a, b**) and 2D optical cross-sections (**c, d**) from the same regions. **e**, Schematic overview of the variational autoencoder (VAE) architecture used for unsupervised learning of spatial patterns. The dataset was split into training (90%) and test (10%) subsets. The VAE comprises an encoder and decoder with dense layers (16 neurons each), optimized using mean squared error (MSE) and Kullback–Leibler (KL) divergence loss functions. MSE minimizes the mean square error during training, ensuring a minimal discrepancy between input and reconstructed output, while the KL loss function encourages the learned latent space distribution to closely resemble a standard normal distribution. In the test phase, the test set is utilized to generate the latent space representation (**i**), and validate the reconstruction output (**j**), ensuring the model’s efficacy and accuracy with a mean square error of 0.0665. **f**,**g,** The original coordinates for CXCL12+ cells (purple) from the selected regions R3 and R5 (Fig. 7h) were retrieved and imported back to the original scans (via Imaris) to visualize the local environment of the cells in sinusoid-enriched regions (**f**) and trabecular sinusoid composite (**g**).

**Extended Data Figure 11.**
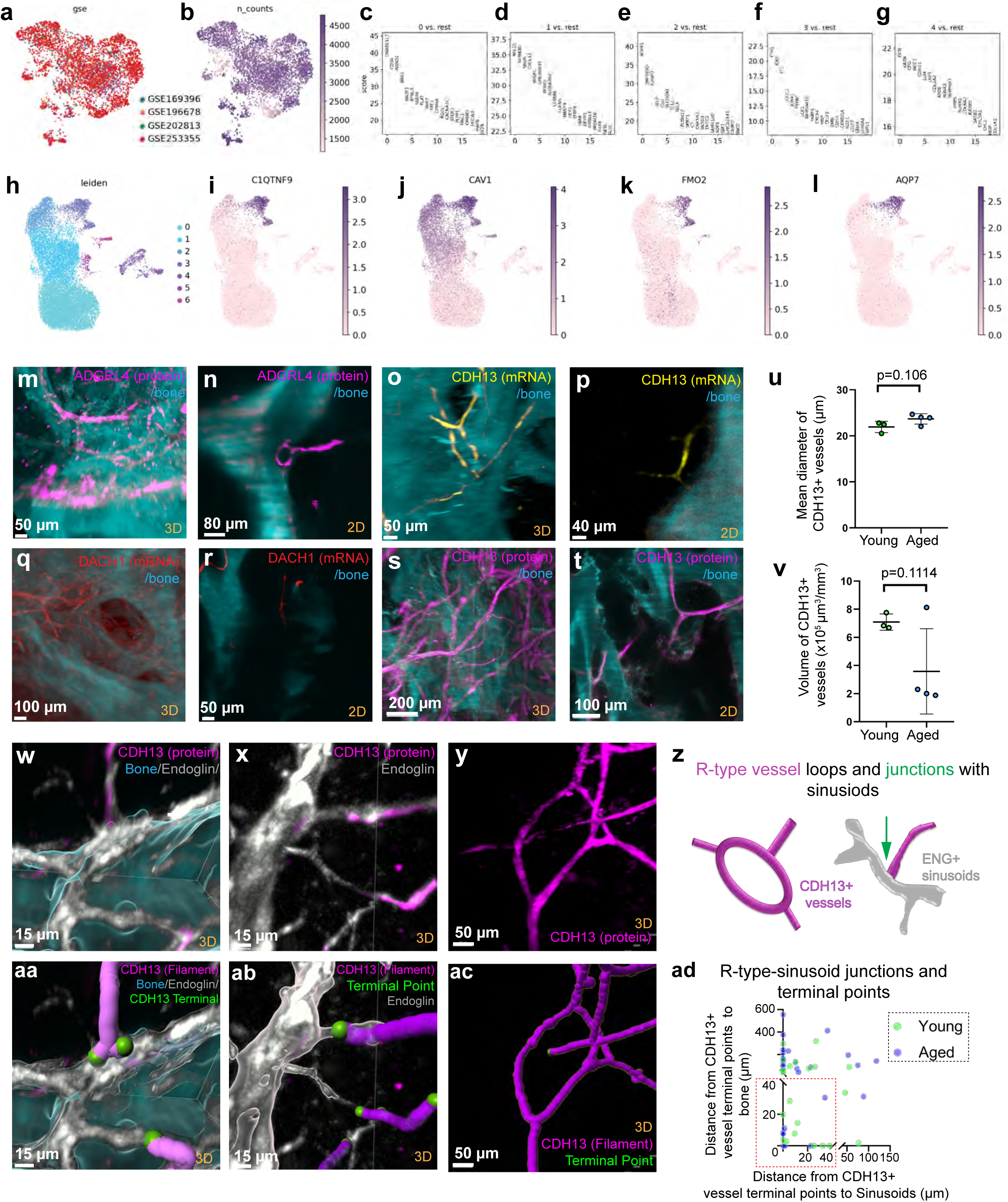
Identification and characterization of R-type vessels near the osteo-sinusoidal niche. **a–g**, Identification of candidate markers for human R-type vessels based on integrated scRNA-seq datasets. **a**, UMAP embedding of pooled human bone marrow endothelial cells (ECs) from our dataset and published sources. **b**, Feature plot showing the number of detected genes per cell. **c–g**, Differential gene expression ranking for each cluster from Main Fig. 8a. **h–l**, UMAP embedding (**h**) and marker gene expression plots (**i–l**) for mouse ECs derived from published scRNA-seq data, including reported R-type EC markers: C1qtnf9 (**i**), Cav1 (**j**), Fmo2 (**k**), and Aqp7 (**l**). **m–t**, Experimental screening of R-type vessel markers using HCR RNA-FISH and immunofluorescence. Representative 3D projections and corresponding 2D optical sections from human bone biopsies stained for ADGRL4 protein (purple; **m, n**), CDH13 mRNA (yellow; **o, p**), DACH1 mRNA (red; **q, r**), and CDH13 protein (purple; **s, t**). **u–v**, Quantification of CDH13+ vessel properties using filament tracing in Imaris. **u**, Mean vessel diameter. **v**, Total vessel volume normalized to tissue volume. Data represent individual patients (young, green dots, n=3; aged, blue dots, n=4). Statistical analysis: unpaired t-test. **w–ad**, Structural features and spatial relationships of CDH13+ R-type vessels with endoglin+ (ENG+) sinusoids and bone. Representative 3D scans showing CDH13 (purple), ENG (white), and bone (cyan, 405 nm autofluorescence) in regions distal (**w**) and proximal (**x**) to trabecular bone. Segmented renderings and terminal points of CDH13+ vessels are shown in **aa–ab**. **y–z**, Example of a self-looping CDH13+ capillary (**y**) with corresponding segmentation (**ac**) and schematic summary (**z**). **ad**, Scatterplot of terminal points from CDH13+ filaments, showing the shortest 3D distances to both ENG+ sinusoids and the bone surface. The red dashed box denotes termini in dual proximity to both structures—potential junction sites within osteo-sinusoidal composites.

